# tRNA-derived fragments elevated in Alzheimer’s disease promote Tau aggregation

**DOI:** 10.64898/2026.05.17.725757

**Authors:** Ami Kobayashi, Chung-Ta Han, Lien D. Nguyen, Karen Tsay, Mariana BMS Martins, Prakash Kharel, Yanhong Zhang, M. Inmaculada Barrasa, Allison M. Williams, Nana Kunii, Yasutoshi Akiyama, Vikram Khurana, Bradley T. Hyman, Pavel Ivanov, Songi Han, Anna M. Krichevsky

## Abstract

Tauopathies, including Alzheimer’s disease (AD), are driven by pathological Tau aggregation, a process that requires co-factors. Small RNAs (sRNA) have been proposed as such co-factors, yet little is known about endogenous transcripts that promote Tau pathology. We identify stress-induced tRNA-derived RNAs or fragments (tDRs/tRFs) as the most dysregulated sRNA class in human AD brains, PS19 mice overexpressing mutant human Tau, and human neuronal tauopathy models. Notably, the highly accumulating 5’Glu^CTC^ and 5’Gly^GCC^ tRFs directly bind Tau and induce its phosphorylation, oligomerization, fibril formation, and impact neurite growth. 5’Glu^CTC^ is enriched in pathological Tau precipitates and co-localizes with oligomeric Tau in PS19 mouse brains. Inhibiting 5’Glu^CTC^ mitigates Tau pathology. Furthermore, these tRFs are highly secreted by neurons and can be taken up by recipient cells, contributing to Tau pathology. Our findings establish 5’Glu^CTC^ as a key regulator of Tau aggregation and suggest its inhibition as a promising therapeutic strategy for tauopathies.

**Highlights:**

- Specific stress-induced 5’tRFs strongly accumulate in Alzheimer’s disease brains
- 5’Glu^CTC^ binds and co-localizes with pathologic Tau in neurons and in the brain
- 5’Glu^CTC^ promotes pathologic Tau species while its inhibition reverses these effects
- Specific 5’tRFs are horizontally transferred between neurons

## Introduction

The microtubule-associated protein Tau stabilizes microtubules under physiological conditions. In response to stress and diseases, Tau undergoes phosphorylation, oligomerization, and somatodendritic accumulation^1^. Fibrillar Tau aggregates are a defining hallmark of tauopathies, including Alzheimer’s disease (AD), Frontotemporal dementia (FTD), and Progressive Supranuclear Palsy (PSP), whereas normal neurons lack Tau fibrils^2,3^. Extensive evidence indicates that Tau oligomers and aggregates are central drivers of toxicity in cellular and animal models. Importantly, Tau burden consistently correlates with clinical symptoms and disease progression across tauopathies^6–8^. Tau aggregates can also be transmitted through inoculation in cellular and murine models^2–5^, and Tau reduction is neuroprotective in mouse models ^9–12^. Thus, elucidation the mechanisms underlying Tau aggregation and toxicity is essential for developing targeted therapies.

Tau aggregation requires co-factors^13,14^; however, the requirement for such co-factors in *in vivo* Tau aggregation and their precise roles remain to be defined. While heparin, polyunsaturated fatty acids, and polyphosphates induce Tau aggregation *in vitro*, the resulting structures differ structurally from those observed in tauopathies. RNA, the most abundant cellular polyanion and a known component of Tau aggregates in tauopathies^15–17^, represents the most plausible physiologic co-factor. While Tau-RNA interactions have been proposed to promote Tau aggregation *in vivo*^17–19^, the endogenous RNA species regulating Tau aggregation, neurofibrillary tangles (NFTs), and human brain pathophysiology remain unknown.

tRNA-derived RNAs (tDRs), also termed tRNA fragments (tRFs)^20^, are abundant small RNAs (sRNAs) implicated in diverse cellular processes^21–27^. Stress-induced tRFs arise from angiogenin (ANG)-mediated cleavage of mature tRNAs^28^ and are classified as 5′tRFs, 3′tRFs, or internal tRFs, with thousands generated from ∼400 human tRNA genes^29^. In proliferating cells, tRFs regulate translation, mRNA silencing, stress responses, and apoptosis^21,23–25,30,31^. While little is known about the role of tRFs in neurons, increased production of ANG-dependent tRFs, particularly 5′tRFs, has been reported in aging mouse brain and in AD hippocampus^32,33^. tRFs are also highly enriched among extracellular RNAs (exRNA) and implicated in intercellular communication^34,35^.

Here, we show that specific 5’tRFs derived from tRNA^Glu-CTC^ (5’Glu^CTC^) and tRNA^Gly-GCC^ (5’Gly^GCC^) are elevated in AD cortex, stressed neurons, and in the hippocampus of Tau-mutant PS19 mice. These tRFs bind Tau, promote its phosphorylation and aggregation, and contribute to neuronal dysfunction. Inhibition of 5’Glu^CTC^ reduces Tau phosphorylation and oligomerization and improves neurite branching and maturation. We further demonstrate that stressed neurons release high levels of 5’Glu^CTC^ and 5’Gly^GCC^ into conditioned media, where they are taken up by recipient neurons. Horizontal transfer of these 5’tRFs is associated with Tau aggregation in recipient cells. Together, these findings identify specific 5’tRFs as endogenous promoters of Tau aggregation and fibril formation and reveal a potential therapeutic target for Tau-associated neurodegenerative disorders.

## Results

### Small RNA analysis identifies tRFs upregulated in human AD patients’ brain cortex

The sRNA profile of normal human, non-pathological temporal cortex shows several distinct bands, with the shortest prominent band (∼75-95nt) corresponding to full-length tRNAs (fl-tRNAs). In contrast, age-matched post-mortem temporal cortex samples from Braak III (mild cognitive impairment) and Braak VI (Alzheimer’s disease, AD) brains exhibited an additional robust ∼30-nt peak, consistent with tRFs (**Fig. 1a**).

**Fig. 1.**
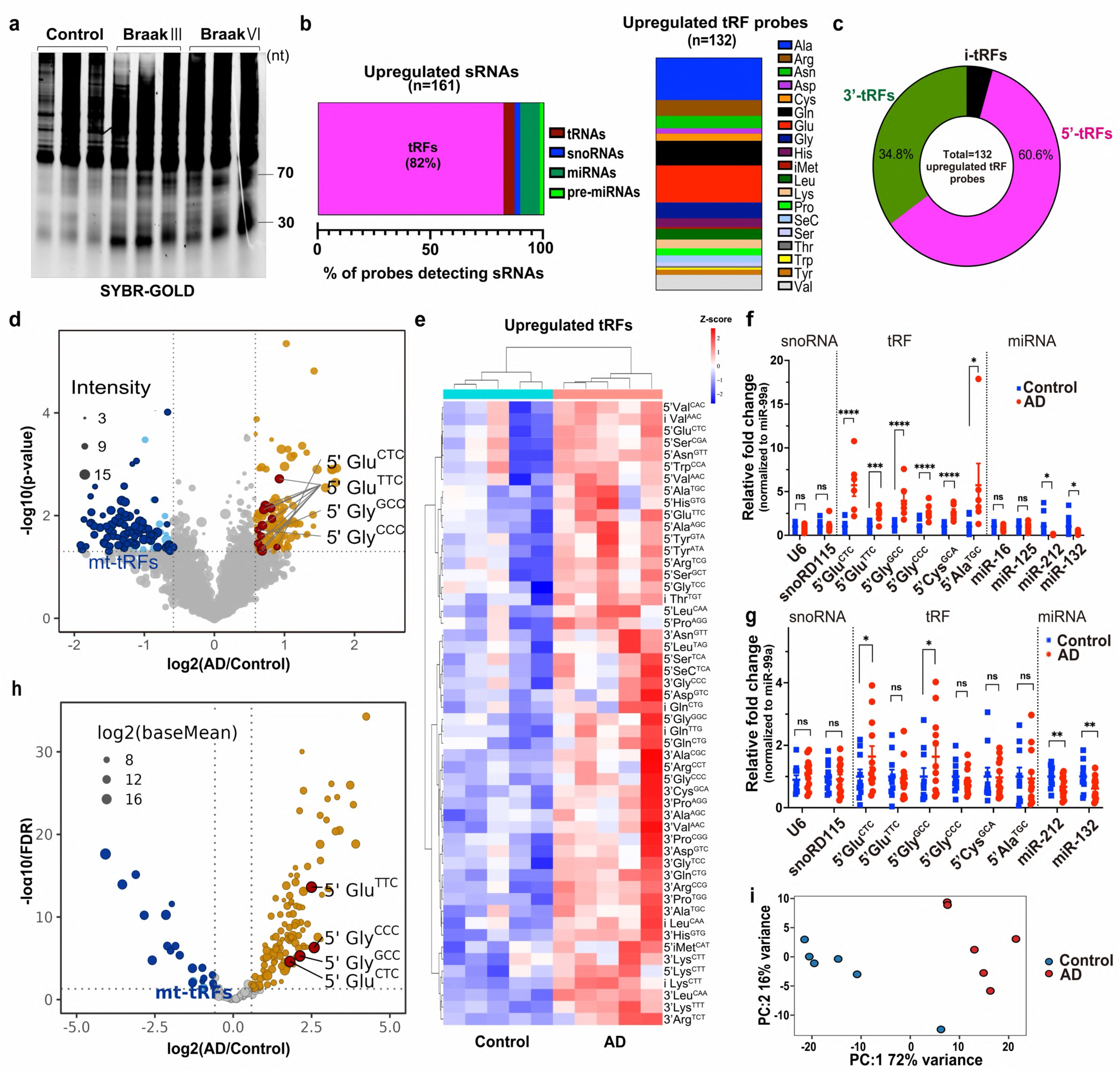
Specific tRFs are upregulated in the AD cortex. **a.** SYBR Gold–stained 15% TBE–urea gel of temporal cortex total RNA from control, Braak stage 3, and Braak stage 6 donors (n = 3/group). **b.** Probe composition of 161 upregulated sRNAs (left) and isotype distribution of 132 upregulated tRF probes (right) in AD temporal cortex. **c**. Overview of tRFs upregulated in AD temporal cortex. **d**. sRNA array volcano plot comparing control (Braak 0) and AD (Braak 6) temporal cortex (n = 5/group). Significantly downregulated and upregulated tRFs are shown in blue and brown, respectively. Dot size indicates signal intensity, and the most abundant 5′tRFs are highlighted in red. **e.** Heatmap of upregulated tRFs in control and AD temporal cortex. **f,g**. qRT-PCR validation of selected tRFs, snoRNAs, and miRNAs in independent New York Brain Bank (**f**; n = 5/group), and Mount Sinai Brain Bank (**g**; n = 6-10/group) cohorts normalized to miR-99a. **h**. Volcano plot of reanalyzed prefrontal cortex sRNA-seq data (GSE48552; control Braak 0–1 vs. AD Braak 5–6; n = 6/group). Significantly downregulated and upregulated tRFs are shown in blue and brown, respectively. tRF counts were summed by isodecoder for DESeq2 analysis; dot size indicates log₂ mean expression, and the most abundant 5’tRFs are highlighted in red. **i.** PCA of tRF profiles in control and AD samples. Unpaired two-tailed Student’s *t-test* was performed in (**f-g**). DESeq2 with Benjamini–Hochberg correction was used in (**h**). Data are mean ± SEM. ****p < 0.0001, ***p < 0.001, **p < 0.01, *p < 0.05, ns = not significant.

To define sRNAs dysregulated in AD, we profiled gray matter of the temporal cortex from Braak VI AD patients and age-matched neurologically normal controls using an sRNA array platform (**Table S1a,b**). Although different sRNA classes cannot be quantitatively assessed by a single RNA-seq strategy due to size and modification differences, the array platform enables such assessments. The array included 9,446 probes, of which 6,287 detected sRNAs: 3,534 tRFs, 133 fl-tRNAs, 407 pre-miRNAs, 1,931 miRNAs, and 282 snoRNAs (**Supplementary Data 1)**. Robust detection fl-tRNAs and tRFs showed separation of expression profiles of the two RNA classes across samples on the principal component analysis (PCA) (**Fig. S1a**). 329 probes detected sRNAs dysregulated in AD (p-value < 0.05, fold change ≥1.5) (**Supplementary Data 1**), including 161 upregulated species, of which 132 (82%) were tRFs (**Fig. 1b; Supplementary Data 1)**. tRFs were the most prominently upregulated class, with several already abundant tRFs further increased in AD (**Fig. S1b,c**). These tRFs derived from 19 tRNA isotypes, with alanine, glutamate, glutamine, glycine, valine, and arginine representing the top six (**Fig. 1b**). Mapping of tRF boundaries revealed enrichment around the anticodon stem–loop, consistent with ANG-mediated cleavage (**Fig. S1d**)^21,28,36,37^. Notably, 80 of 132 upregulated tRF probes (60.6%) corresponded to 5′tRFs (**Fig. 1c)**.

Among these, 5′Glu-derived tRFs were most prominent: 19 probes detected 5’Glu^CTC^ and 5’Glu^TTC^ tRFs (**Fig. 1d; Supplementary Data 2,3**). Highly abundant baseline tRFs, 5’Glu^CTC^, 5’Glu^TTC^, 5’Gly^CCC^, and 5’Gly^GCC^, were all significantly elevated in AD cortex (**Fig. 1d**). Overall, AD samples displayed a distinct signature of 52 upregulated tRFs (**Fig. 1e**). In contrast, most downregulated tRFs were mitochondrial-derived (mt-tRFs): 80.2% of downregulated probes corresponded to mt-tRFs, with 77.9% being 5′mt-tRFs (**Fig. S1d–f; Supplementary Data 4**).

Several AD-elevated 5′tRFs belong to the highly abundant sRNAs in the primate brain^38–40^. To validate the results, we measured the expression of selected sRNAs (one snoRNA, six tRFs, and five miRNAs) in two cohorts of temporal cortex samples using sRNA-specific qRT-PCR, confirming increased levels of 5’Glu^CTC^ and 5’Gly^GCC^ in both AD cohorts (**Fig. 1f,g**). As expected, miR-132, an miRNA linked to both amyloid and Tau pathology^41–47^, was downregulated (**Fig. S1g)**. Reanalysis of an independent sRNA-seq dataset from AD prefrontal cortex (GSE48552)^48^ confirmed widespread tRF accumulation, with 5’Glu^CTC^ and 5’Gly^GCC^ being the most abundant and further elevated in AD **(Fig. 1h; Fig. S1h; Supplementary Data 5)**. PCA based on all 368 tRF isodecoders clearly separated AD and control samples **(Fig. 1i)**.

To assess the specificity of 5’Glu^CTC^ and 5’Gly^GCC^ accumulation in AD, we reanalyzed published sRNA datasets from Frontotemporal Lobar Dementia with Tau pathology (FTLD-MAPT; E-MTAB-12731, batch 1)^49^ and sporadic Parkinson’s Disease (PD; GSE72962, GSE 64977)^50,51^ (**Supplementary Data 6,7)**. 5’Glu^CTC^ and 5’Gly^GCC^, together with another highly abundant 5’Gly^CCC^, were upregulated in the prefrontal cortex of FTLD-MAPT, mirroring the pattern observed in AD (**Fig. S2a-d)**. In contrast, these 5’tRFs were not elevated in the PD prefrontal cortex (**Fig. S2e-h)**, indicating their association with tau pathology rather than a general neurodegenerative stress response. Overall, these results identify 5’Glu^CTC^ and 5’Gly^GCC^ as the sRNAs commonly accumulated in the cortical brain regions of AD and other tauopathies.

### Chronic neuronal stress upregulates 5’Glu^CTC^ and 5’Gly^GCC^ and increases oligomeric and pTau in rodent and human neurons

tRFs are generated by tRNA cleavage by specific nucleases in response to stress^28,52,53^, and AD is a stress-associated disease^54^. Chronic neuronal stress promotes neuroinflammation and oxidative stress, major contributors to Tau hyperphosphorylation and aggregation^55,56^. To test whether neuronal stress triggers the tRF accumulation observed in AD cortex, we exposed rat primary cortical neurons and human iPSC-derived excitatory neurons (iNs) to low-dose, subtoxic, AD-relevant stressors: glutamate (excitotoxic stress), hydrogen peroxide (oxidative stress), and partially aggregated Aβ1–42 oligomers (1/2t_max_Aβ)^57–60^. Neurotoxicity was dose-dependent (**Fig. S3a,b**), and at selected concentrations (glutamate: 3 µM rat, 0.5 µM human; H₂O₂: 0.5 µM rat, 5 nM human; 1/2t_max_Aβ: 0.25 µM rat, 0.5 nM human), viability loss did not exceed 10% at 24h (**Fig. S3a,b**). No morphological abnormalities or increased apoptosis were detected by imaging or 7-AAD/ Annexin V assays (**Fig. 2a; Fig. S3c**).

**Fig. 2.**
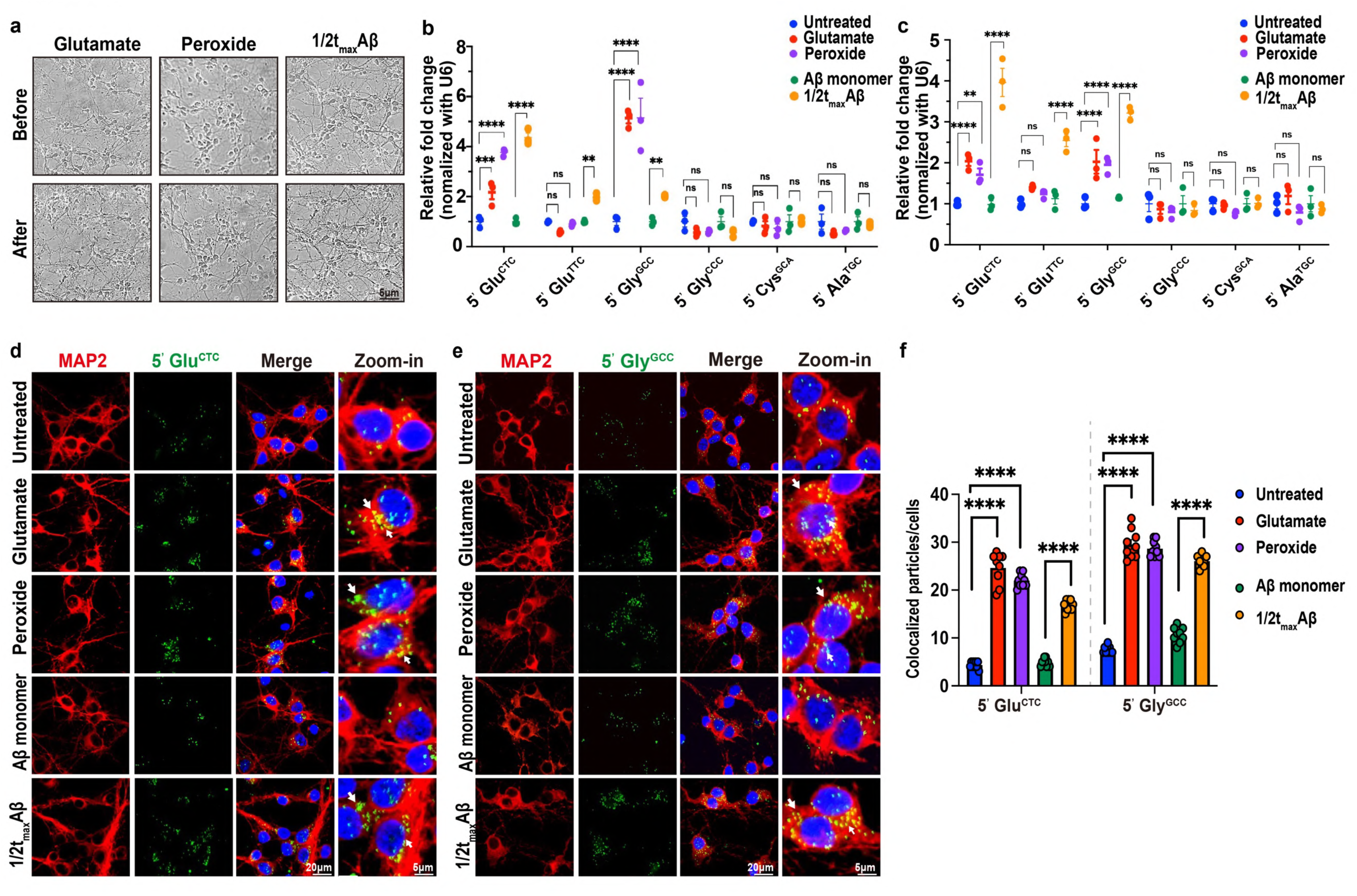
Specific tRFs are upregulated in neurons under persistent extrinsic stress. **a.** Representative bright-field images of rat neurons before and 24h after treatment with subtoxic doses of 3 µM glutamate, 0.5 µM peroxide, and 0.25 µM 1/2t_max_Aβ. **b,c.** qRT-PCR analyses of tRFs in rat (**b**) and human (**c**) neurons, normalized to U6 (n = 3/group). **d,e.** Representative FISH/IF images of (**d**) 5’Glu^CTC^ or (**e**) 5’Gly^GCC^ FISH (green) in unstressed and stressed rat neurons. MAP2 IF (red) and DAPI-stained nuclei (blue). **f.** Quantification of 5’tRF puncta in MAP2^+^ cells (n = 3 independent neuronal cultures; ≥200 cells/condition). One-way ANOVA with Dunnett’s test was performed in (**b-c**) and (**f**). All graphs show mean ± SEM. ****p < 0.0001, ***p < 0.001, **p < 0.01, *p < 0.05, ns = not significant.

Despite minimal toxicity, sustained low-level stress selectively upregulated 5’Glu^CTC^ and 5’Gly^GCC^ in both rat and human neurons (**Fig. 2b, c**). 5’Glu^TTC^ was increased only by 1/2t_max_Aβ (**Fig. 2b, c**). To visualize intracellular 5’Glu^CTC^ and 5’Gly^GCC^, we performed fluorescence *in situ* hybridization (FISH). FISH confirmed increased cytoplasmic and perinuclear 5’Glu^CTC^ and 5’Gly^GCC^ signals across all stress paradigms (**Fig. 2d-f**). The signals were abolished by pre-transfections of the corresponding antisense oligonucleotides (ASOs), confirming specificity of 5’tRF detection (**Fig. S4a**). In contrast, probes for other fragments from the same parental tRNAs (3’Glu^CTC^, 3’Gly^GCC^, iGlu^CTC^, iGly^GCC^) exhibited distinct patterns and no stress responsiveness **(Fig. S4b**). In stressed neurons, both 5’Glu^CTC^ and 5’Gly^GCC^ showed minimal colocalization with the stress granule marker G3BP1^61,62^ **(Fig. S4c**).Because glutamatergic, oxidative, and Aβ stresses promote Tau hyperphosphorylation^63–66^, we assessed Tau pS396 (associated with early disease stage) and Tau oligomerization using Tau 22 and TOMA1 antibodies that recognize misfolded oligomeric Tau (**Fig. S5a**). Western blot (WB) showed induction of phosphorylated and oligomeric, but not total, Tau in stressed rat and human neurons (**Fig. S5b,c**). Consistent with these results, increased cytoplasmic oligomeric Tau puncta was observed by immunofluorescence (**Fig. S5d**).

### 5’GluCTC and 5’Gly^GCC^ are upregulated by intrinsic Tau-induced stress in human neurons and in the mouse brain

Tau mutations are major genetic drivers of Tau pathology, disrupting cellular homeostasis, transport, and synaptic function through toxic gain-of-function effects. Although AD itself lacks Tau mutations, the PS19 mouse model overexpressing human 4R1N P301S Tau is a widely used and robust model of ADRD neurodegeneration^67^. To assess whether intrinsic Tau-related stress alters tRF levels, we analyzed hippocampi from WT and PS19 mice. qRT-PCR revealed increased 5’Glu^CTC^, 5’Gly^GCC^, 5’Gly^CCC^, and 5’Cys^GCA^ as early as 4 months (presymptomatic), with sustained elevation at 10 months (**Fig. 3a,b**). Northern blotting (NB) confirmed strong accumulation of 5’Glu^CTC^ and 5’Gly^GCC^ in PS19 hippocampi (**Fig. 3c**); the larger fold-changes (up to 7–18 fold) likely reflect detection of closely related 30–34-nt tRFs by NB, whereas qRT-PCR is isoform-specific. Corresponding full-length tRNAs were unchanged (**Fig. 3c**). FISH of the CA1 region corroborated 5’Glu^CTC^ and 5’Gly^GCC^ accumulation, with perinuclear speckles across layers and strongest enrichment in the pyramidal layer (**Fig. 3d,e**).

**Fig. 3.**
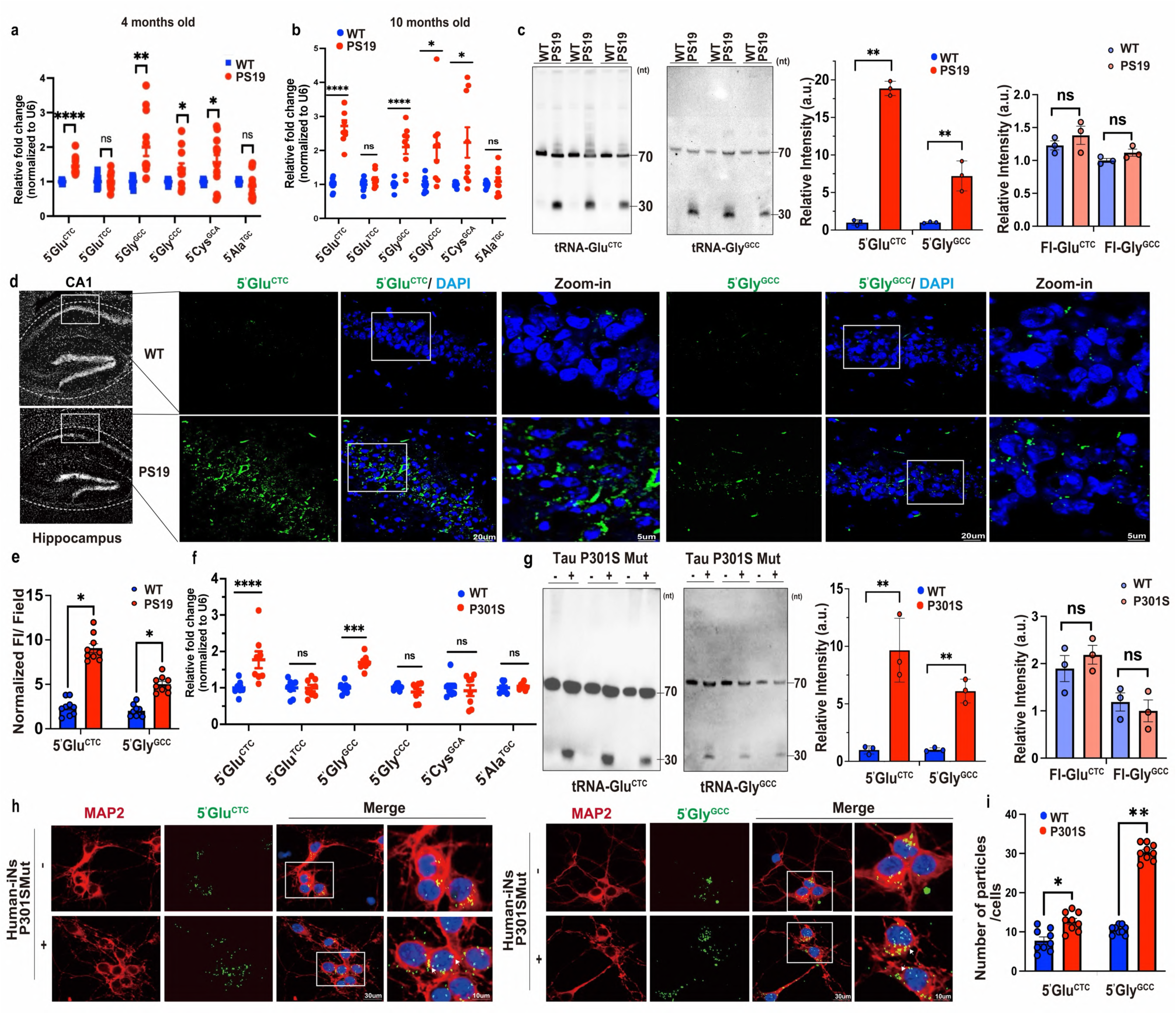
5’Glu^CTC^ and 5’Gly^GCC^ are upregulated in PS19 mice and Tau P301S_Mut_ human neurons. **a,b.** qRT-PCR analysis of indicated tRFs in hippocampi from WT and PS19 mice at 4 months (**a**) and 10 months (**b**), normalized to U6 snRNA. n = 9 mice/group. **c.** Northern blot and quantification of 5’Glu^CTC^, Gly^GCC^, and corresponding full-length tRNAs in 10-month-old WT and PS19 mice hippocampi (n = 3/group). **d,e.** Representative hippocampal FISH images (**d**) and quantification (**e**) 5’Glu^CTC^ and 5’Gly^GCC^ (green) in 10-month-old WT and PS19 mice. Nuclei (blue). Inset show CA1 (n = 9 sections from 3 mice). **f.** qRT-PCR analysis of indicated tRFs in WT and Tau P301S_Mut_ human neurons, normalized to U6 snRNA (n = 8/group). **g.** Northern blot and quantification of 5’Glu^CTC^ and 5’Gly^GCC^ in WT and Tau P301S_Mut_ human neurons (n = 3/group). **h,i.** Representative images of 5’Glu^CTC^ or 5’Gly^GCC^ FISH (green), MAP2 IF (red), nuclei (blue) (**h**) and quantification of 5’Glu^CTC^ and 5’Gly^GCC^ puncta in WT and Tau P301S_Mut_ human neurons (**i**). (n = 3/group; ≥300 cells/condition). Unpaired two-tailed Student’s *t-test* was performed in (**a-c**)(**e-g**)(**i**). Data are mean ± SEM.: ****p < 0.0001, ***p < 0.001, **p < 0.01, *p < 0.05, ns = not significant.

To extend these findings to human neurons, we analyzed iNs carrying an autosomal-dominant Tau P301S mutation (P301S_Mut_), and the corrected WT line. qRT-PCR and NB showed upregulation of 5’Glu^CTC^ and 5’Gly^GCC^ in P301S_Mut_ neurons (**Fig. 3f,g**), and FISH confirmed their cytoplasmic and perinuclear accumulation (**Fig. 3h,i**). Together, these data indicate that intrinsic Tau stress from P301S mutation, like extrinsic stressors, selectively elevates specific 5′tRFs, implicating them in Tau-related pathology.

### 5’GluCTC is enriched in Tau complexes in rodent and human neurons under intrinsic and extrinsic stress

NFTs isolated from AD brains contain RNA^17,68,69^, and Tau-RNA interactions may promote Tau aggregation^17,68^. sRNA, and particularly tRNAs, have emerged as candidate species^17^. However, whether Tau binds fl-tRNAs or tRFs, and whether such interactions are selective, remains unknown. Because fl-tRNAs are highly structured and engaged in translation, we hypothesized that less structured tRFs that accumulate during stress and in AD are the predominant endogenous Tau binders.

To test this, we performed 5′tRF pull-down assays in neurons transfected with synthetic biotinylated 5′tRF mimics, including 5’Glu^CTC^ and 5’Gly^GCC^, designed with similar end-modifications to control for sequence-independent effects (**Table S2**). Following cross-linking and stringent streptavidin-based purification (**Fig. 4a**), WB analysis revealed selective co-precipitation of phosphorylated (pS396) and oligomeric (Tau22) Tau with 5’Glu^CTC^ and 5’Gly^GCC^, but not with other 5′tRFs (**Fig. 4b, Fig. S6a**), indicating specific 5’tRF interactions with pathological Tau species.

**Fig. 4.**
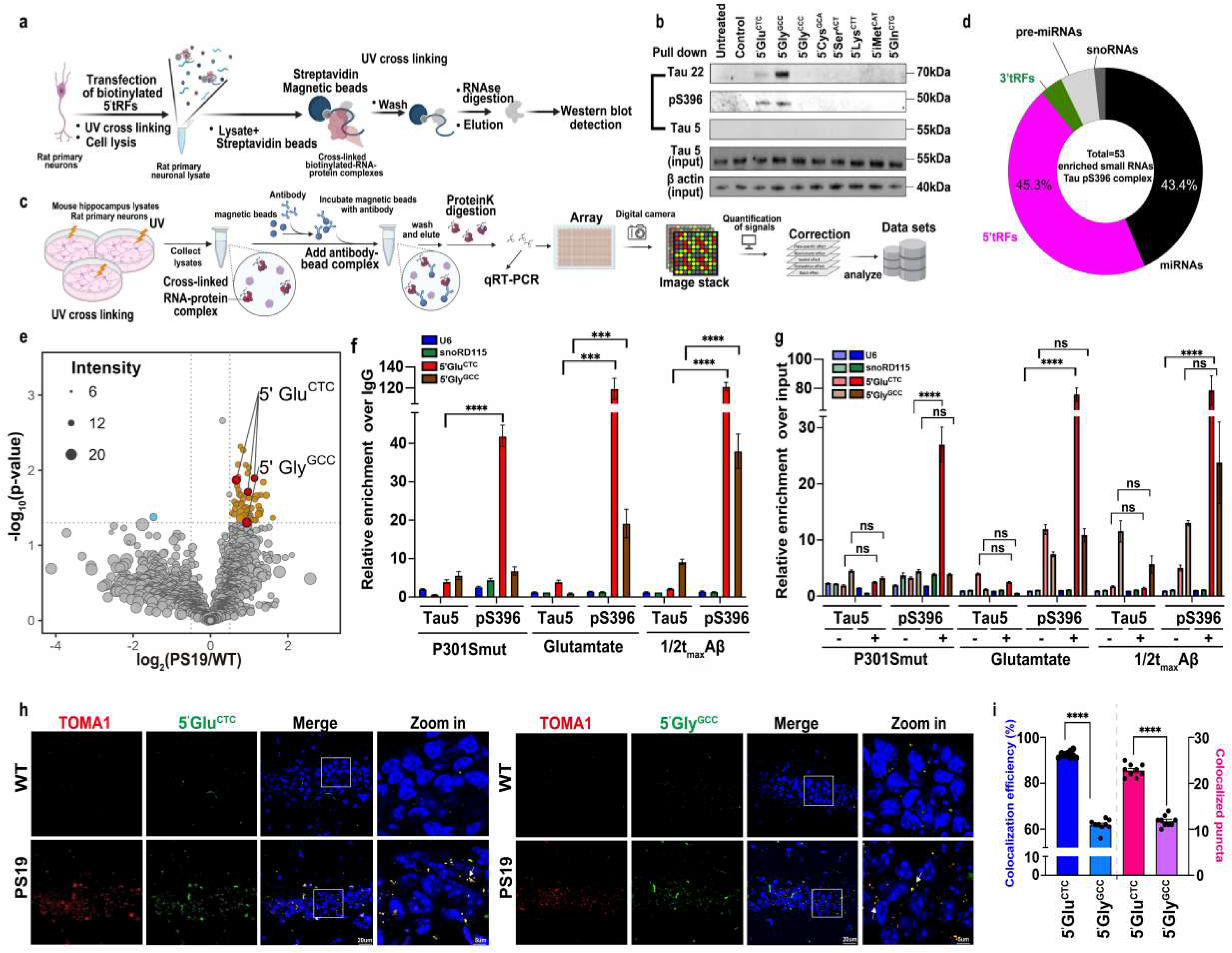
Specific 5’tRFs selectively bind to pathological Tau in stressed neurons and mouse PS19 hippocampus. **a.** Schematic of the 5′tRF pull-down assay. **b.** Biotin pull-down in rat primary neurons transfected with biotinylated 5’tRFs. Representative WB show selective binding of 5’Glu^CTC^ and 5’Gly^GCC^ to oligomeric Tau (Tau 22), pTau (Tau pS396), and total Tau (Tau 5). **c.** Schematic of the modified CLIP experiment. **d.** sRNA classes among 53 RNAs enriched in pS396-Tau immunoprecipitates. **e.** CLIP-sRNA array volcano plot from WT and PS19 mice hippocampi immunoprecipitated with Tau pS396 antibody. Significantly downregulated and upregulated tRFs are shown in blue and brown, respectively; dot size indicates signal intensity, and the most abundant 5′tRFs are highlighted in red. (n = 3/group). **f,g.** CLIP-qRT-PCR analysis of selected sRNAs in PS19 mouse hippocampi and stressed rat neurons. Data are shown as fold enrichment in Tau pS396 complexes relative to Tau 5, and normalized to IgG (**f**) and RNA(**g**) input (n = 3/group). **h,i.** Representative CA1 images of 5’Glu^CTC^ or 5’Gly^GCC^ FISH (green), TOMA1 IF (red), nuclei (blue) (**h**) and quantification of TOMA1–5′tRF colocalization (**i**) shown as colocalized puncta count/field (right panel) and colocalization efficiency as a ratio to total 5’tRF puncta/field (left panel) in 10-month-old WT and PS19 mice (n = 3/group; ≥150 cells/condition). Arrows indicate colocalization. One-way ANOVA with Dunnett’s test was performed in (**f**)(**g**) and unpaired two-tailed Student’s *t-test* was performed in (**i**). Data are mean ± SEM. ****p < 0.0001, ***p < 0.001, **p < 0.01, *p < 0.05, ns = not significant.

We next performed modified CLIP in PS19 mouse hippocampus and rat primary neurons using antibodies to phosphorylated (pS396, PHF1) or total Tau (Tau5), followed by sRNA array profiling (**Fig. 4c**). Multiple 5’tRFs were enriched in Tau pS396 complexes from PS19 hippocampi (**Fig. 4d, Supplementary Data 8-10**), with 5’Glu^CTC^ among the most enriched (**Fig. 4e**). 5’Gly^GCC^ showed weaker enrichment. CLIP–qRT-PCR confirmed significant enrichment of 5’Glu^CTC^ in Tau pS396 and PHF1 complexes **(Fig. 4f, Fig. S6b,c)**. It was significant relative to total Tau complexes and the enrichment persisted after normalization to elevated tRF levels in PS19 tissue **(Fig. 4g; Fig. S6b,c)**. In contrast, 5’Glu^CTC^ showed no enrichment in total Tau complexes in both WT and PS19 hippocampi when normalized to tRF levels **(Fig. 4g)**. 5’Gly^GCC^ was not significantly enriched in Tau 5 or Tau pS396 complexes from PS19 hippocampi **(Fig. 4f,g)**. Although a modest enrichment was observed in PHF1 complexes relative to total Tau **(Fig. S6b**), this effect was lost after RNA normalization (**Fig. S6c**). CLIP-qRT-PCR in naive and glutamate- or 1/2t_max_Aβ-stressed primary neurons similarly showed selective enrichment of 5’Glu^CTC^ in Tau pS396 and PHF-1 complexes compared to total Tau complexes, which remained significant after normalization to RNA input **(Fig. 4f,g; Fig. S6b,c)**. Apparent 5’Gly^GCC^ enrichment in Tau pS396 complexes in stressed neurons was abolished by RNA normalization **(Fig. 4f,g)**, and no 5’Gly^GCC^ enrichment in stressed or naïve neurons was observed in PHF-1 complexes **(Fig. S6b,c)**.

We next examined colocalization of 5’Glu^CTC^ and 5’Gly^GCC^ with oligomeric Tau in mouse hippocampus, using combined FISH and immunofluorescence. Whereas minimal tRF and TOMA1 signals and colocalization were observed in 10 m.o. WT mice, PS19 mice showed elevated 5’Glu^CTC^ and 5’Gly^GCC^, with their substantial co-localization with TOMA1 (**Fig. 4h**). About 62% of 5’Gly^GCC^ and 92% of 5’Glu^CTC^ puncta co-localized with TOMA1 (**Fig. 4i**). Imaging of stressed rat and human neurons, as well as Tau P301S_Mut_ human iNs, revealed overlap between 5’Glu^CTC^ (green) and oligomeric Tau (TOMA1, red) intensity peaks (**Fig. S6d**). The overlap between 5’Gly^GCC^ and TOMA1 was limited (**Fig. S6e**). Quantification confirmed increased co-localization of 5’Glu^CTC^ with TOMA1 puncta across all models, whereas 5’Gly^GCC^ showed minimal co-localization (**Fig. S6f,g**).

Together, these complementary approaches across multiple models demonstrate a selective interaction and colocalization of 5’Glu^CTC^ with phosphorylated and oligomeric Tau in stressed and Tau-mutant neurons.

### Specific 5’tRFs induce Tau phosphorylation and oligomerization in neurons

5’Glu^CTC^ is an abundant sRNA in the brain^38^ that further accumulates in AD tissues^32^. To test whether elevated 5’Glu^CTC^ affects Tau metabolism, we overexpressed a synthetic 5’Glu^CTC^ mimic in neurons, alongside mimics of other abundant neuronal tRFs. Notably, 5’Glu^CTC^, 5’Gly^GCC,^ and 5’Gly^CCC^ induced Tau phosphorylation at several residues associated with intraneuronal NFT (S396, Thr181, and S262) and promoted Tau misfolding and oligomerization (Tau22 and TOMA1), without altering total Tau, in both rat primary and human neurons 24 hours after transfection (**Fig. 5a, Fig. S7a**). Phosphorylation at S262 was associated with appearance of oligomeric Tau species. In contrast, these 5’tRFs had no effects on phosphorylation at Thr231, an early pre-NFT epitope (**Fig. 5a**). 5’Cys^GCA^ also increased pS396 and Tau22 signals, whereas other tested 5’tRFs had no effects (**Fig. 5a, Fig. S7a**). Consistently, immunofluorescence revealed a 3-4-folds increase in cytoplasmic TOMA1-positive Tau puncta in neurons overexpressing 5’Glu^CTC^, 5’Gly^GCC^, and 5’Gly^CCC^ (**Fig. 5b,c; Fig. S7b,c)**. X-34 positive β sheet-rich aggregates were observed exclusively in these tRF-overexpressing cultures, and they overlapped with TOMA1 puncta (**Fig. 5b,c**). Furthermore, while no differences were observed in the sarkosyl-soluble Tau fractions isolated from the panel of tRF-overexpressing neuron cultures, 5’Glu^CTC^, 5’Gly^GCC^, and 5’Gly^CCC^ increased the amount of insoluble Tau species, consistent with enhanced Tau fibril formation (**Fig. 5d**).

**Fig. 5.**
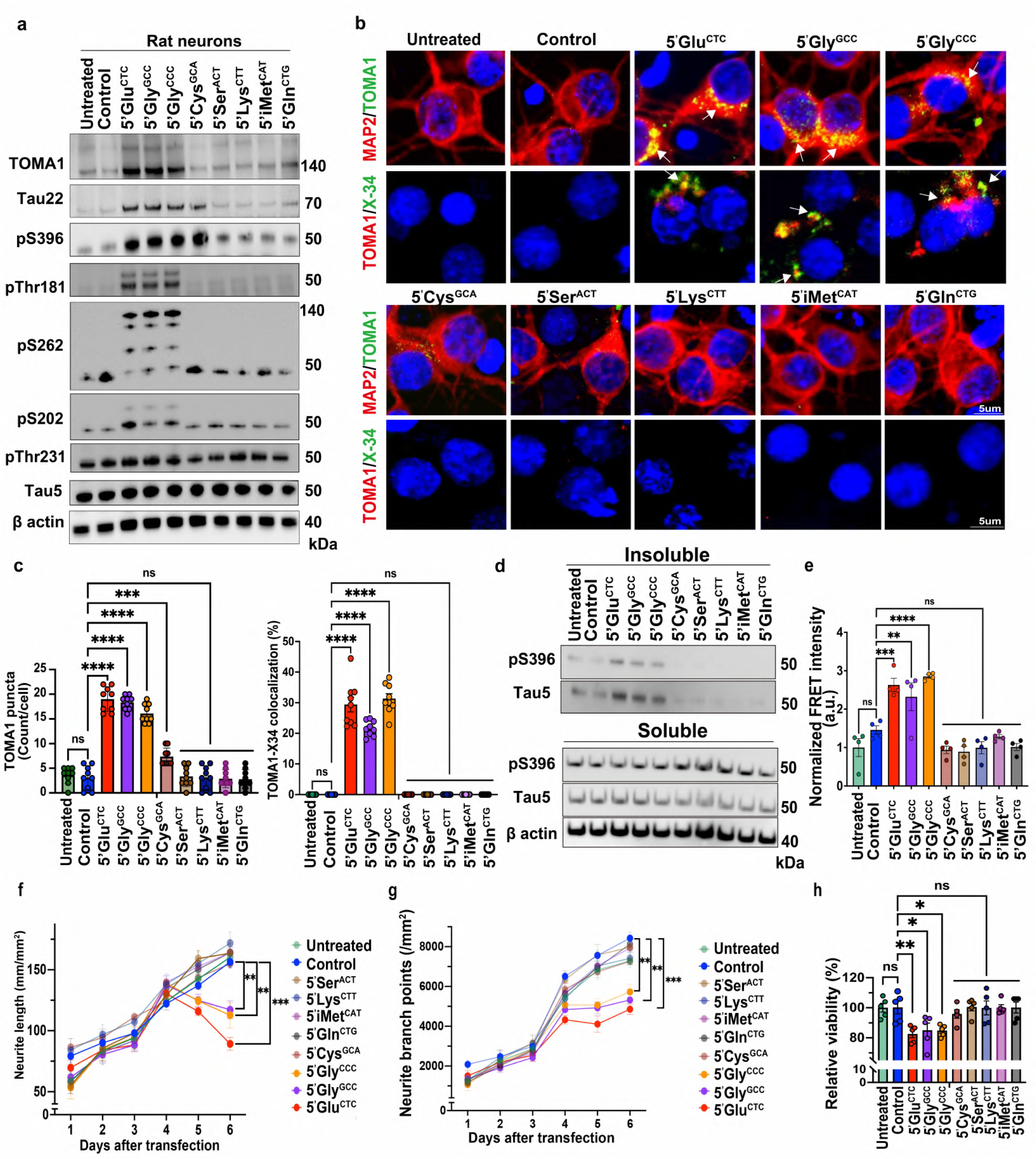
5’Glu^CTC^ and 5’Gly induce Tau phosphorylation, oligomerization, aggregation, and impair neuronal viability. **a.** Representative WB of oligomeric Tau (TOMA1, Tau 22), pTau (pS396, pThr181, pS262, pS202, pThr231), and total Tau (Tau 5) in rat neurons transfected with indicated 5’tRFs along with control cultures (n = 3/group). **b,c.** Representative IF and X-34 staining images (**b**) and quantification (**c**) of TOMA1 puncta and TOMA1/X34 colocalization in 5’tRF-transfected rat neurons. Colocalized area (yellow), and nuclei (blue). Arrows mark TOMA1 puncta colocalized with MAP2 or X-34 (n = 3/group; ≥250 cells/condition). **d.** WB of Tau species in sarkosyl-soluble and -insoluble fractions. **e.** Tau seeding assay in HEK-tau-biosensor cells transfected with indicated 5’tRFs and HMW Tau shown as normalized FRET intensity (n = 4/group). **f,g.** Live-cell imaging quantification of neurite length (**f**) and neurite branch points (**g**) in rat neurons over 6 days after 5′tRFs transfection (n = 3/group). **h.** Neuronal viability 6 days after 5′tRF transfection (n = 5/group). One-way ANOVA with Dunnett’s test was performed in (**c**)(**e**)(**f**)(**g**) and (**h**). Data are mean ± SEM. ****p < 0.0001, ***p < 0.001, **p < 0.01, *p < 0.05, ns = not significant.

We next examined whether these 5’tRFs promote Tau aggregation in tau-biosensor HEK cells seeded with high-molecular-weight Tau from AD brain^70,71^. Remarkably, 5’Glu^CTC^, 5’Gly^GCC^, and 5’Gly^CCC^, but not other tRFs, induced Tau seeding, as measured by normalized FRET intensity (**Fig. 5d**). To assess the effects of 5’tRFs on neuronal morphology and health, we imaged transfected neurons over 6 days and monitored neurite lengths and branching. 5’Glu^CTC^, 5’Gly^GCC^, and 5’Gly^CCC^ impaired neuronal growth, reducing neurite length and branching in both rat and human neurons, compared to untreated cultures and neurons transfected with other tRFs (**Fig. 5e,f, Fig. S7d,e**). Moreover, they reduced neuronal viability by 17-20% (**Fig. 5g, Fig. S7f**). Together, these data indicate that specific 5′tRFs elevated in AD, particularly 5’Glu^CTC^ and 5’Gly^GCC^, compromise neuronal health by inducing Tau phosphorylation and oligomerization.

### 5’GluCTC and 5’Gly^G/CCC^ promote Tau aggregation and fibril formation by directly interacting with specific Tau residues

To further investigate the stand-alone ability of tRFs to bind Tau and promote its aggregation, we employed an *in vitro* Thioflavin T (ThT) assay as a semiquantitative measure of Tau fibril formation. We employed a seeding-competent Tau 295–313 peptide (jR2R3) encompassing the fibrilization-prone PHF6 motif (VQIVYK) (**Fig. 6a**), capable of forming fibrils *in vitro* and inducing aggregation of cellular Tau over multiple generations^72^, with P301L substitution further enhancing fibrillization^73^. This minimal system allowed us to probe whether specific tRFs interact with the PHF6-containing R2–R3 junction, a region critical for Tau aggregation initiation. 5’Glu^CTC^, 5’Gly^GCC^, and 5’Gly^CCC^, but not other tRFs, induced fibrillization of this peptide (**Fig. 6b**). Dense fibrils induced by these tRFs were confirmed by transmission electron microscopy (TEM) (**Fig. 6c**). While these tRFs robustly promoted fibrillization of jR2R3-P301L Tau, they did not induce significant fibril formation of full-length 0N4R Tau, which is stabilized in an aggregation-resistant conformation^74^.

**Fig. 6.**
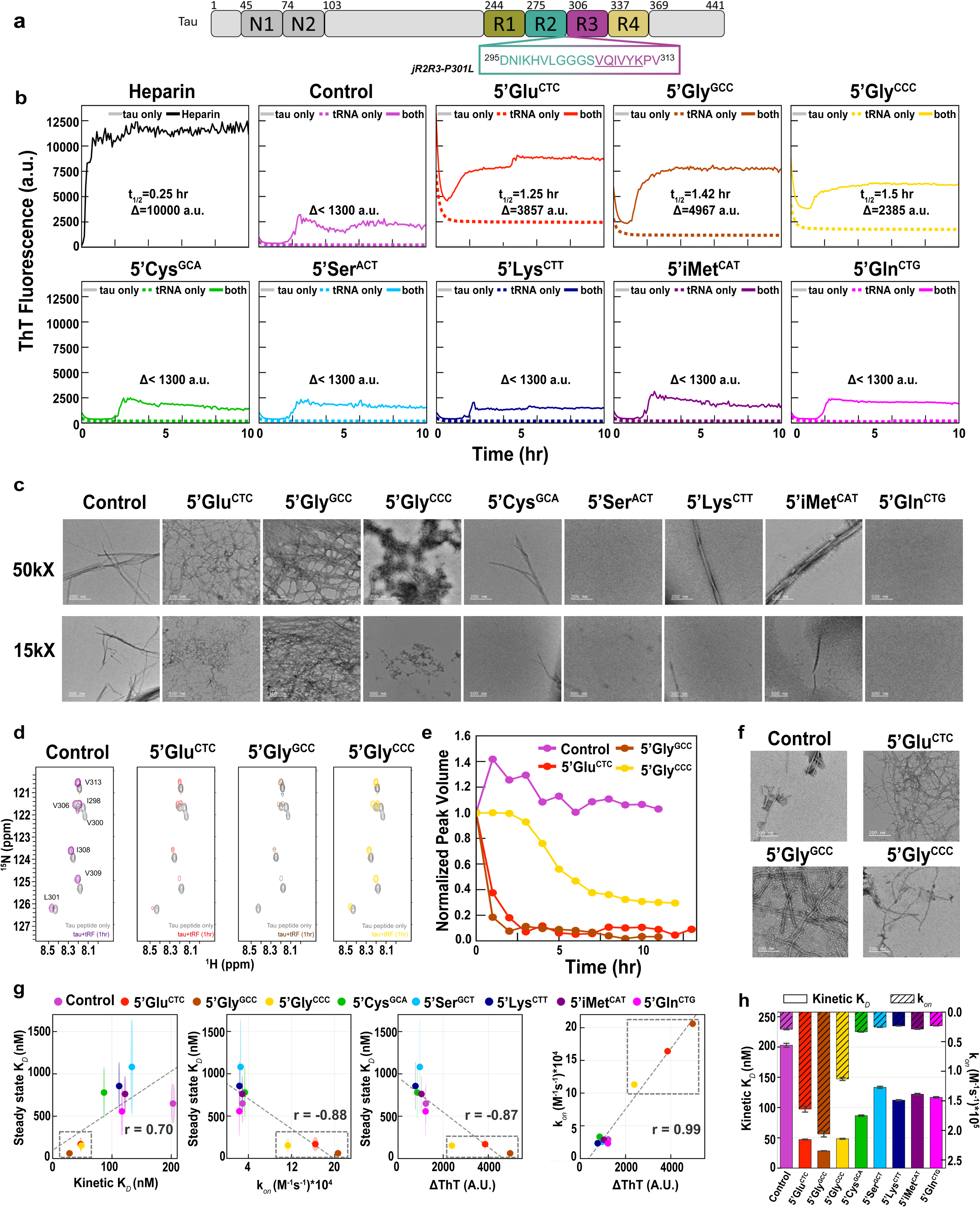
5’Glu^CTC^ and 5’Gly bind Tau-seeding peptide and promote tau fibrilization. **a.** Schematic of Tau, including the jR2R3-P301L peptide at the R2/R3 splice junction, spanning residues D295-V313, with PHF6 sequence underlined. **b.** ThT fluorescence assay of jR2R3-P301L fibrillization with indicated tRFs. Heparin served as a positive control. **c.** Representative TEM images of the generated fibrils. Scale bars: 200 and 500 nm for upper-row and lower-row images, respectively. **d.** NMR analysis of jR2R3-P301L tau peptide in the absence or presence of tRFs. Grey and colored circles denote peaks from seven hydrophobic, isotope-enriched residues before and after tRF addition, respectively. **e.** Time course of normalized NMR peak intensities following tRF addition. **f.** Representative TEM images of fibrils formed in NMR tubes. Scale bars are 200 nm. **g.** Pairwise correlations for steady state *K_D_*, kinetic *K_D_*, association rate constant *k_on_* measured by BLI, and ThT fluorescence enhancement (ΔThT) across nine tRFs. Pearson r values are shown. **h.** Comparison of *K_D_* (solid, lower bars) and association rate constant *k_on_* (hatched, upper bars) across Tau-binding tRFs.

To track Tau monomer – tRF interactions at the molecular level, we employed solution state nuclear magnetic resonance (NMR) sensitive to residue-specific structural changes in molecular dynamics of soluble proteins. We compared ^1^H-^15^N HMQC spectra for jR2R3-P301L peptide that includes 7 ^15^N-isotope-enriched residues before and after addition of 5’Glu^CTC^, 5’Gly^GCC^, 5’Gly^CCC^, alongside control RNA. The tRFs induced peak broadening (i.e., diffusion or disappearance) indicative of fibril formation, with a faster and more extensive changes triggered by 5’Glu^CTC^ and 5’Gly^GCC^ (**Fig. 6d,e**). Consistent with previous observations of jR2R3-P301L forming a strand-loop-strand β-sheet with a hotspot around residue 300^74^, the tRFs induced Tau oligomer formation with residue 300 being immobilized first (**Fig. 6d**). Following the NMR studies, TEM confirmed Tau fibril formation, with the denser fibrils induced by 5’Glu^CTC^ and 5’Gly^GCC^ followed by 5’Gly^CCC^, and no fibrils formed in control conditions (**Fig. 6f**). To assess jR2R3 binding affinity to tRFs, we used Bio-layer interferometry (BLI) and determined *K_D_* values by both unimodal 1:1 Langmuir global kinetic fitting and steady state analysis. The two approaches showed strong concordance across all tRFs (Pearson r = 0.70), indicating a consistent binding hierarchy (**Fig. 6g**). *K_D_* correlated negatively with the association rate constant *k_on_* (Pearson *r* = −0.88), consistent with faster-binding tRFs exhibiting higher affinity. Notably, *k_on_* also correlated strongly with ThT fluorescence enhancement (*r* = −0.99), suggesting that rapid and tight binding induces fibrillization (**Fig. 6g**). Together, these results define relative binding affinities across tRFs, supporting *K_D_* values of 48 nM and 28 nM for 5’Glu^CTC^ and 5’Gly^GCC^, respectively (**Fig. 6h**), and high selectivity of the tRF-Tau interaction. Collectively, these experiments indicate that 5’Glu^CTC^, 5’Gly^GCC^, and 5’Gly^CCC^ directly interact with Tau monomers around residues 300–310 and enhance oligomerization and fibrilization.

### Inhibition of 5’Glu^CTC^ in stressed and Tau-mutant neurons reduces Tau oligomerization, phosphorylation, and enhances neuronal health

Given the accumulation of 5’Glu^CTC^ and 5’Gly^GCC^ in AD and their ability to promote Tau phosphorylation and oligomerization, we tested the effects of tRF inhibitors. We utilized specific ASOs (**Table S3**), optimized for high affinity, potency, specificity, and low-toxicity *in vitro* ^75,76^. Neurons were transfected with tRF-targeting or non-targeting control oligonucleotides (**Table S3**). The anti-tRFs ASOs robustly reduced the cognate 5’Glu^CTC^ and 5’Gly^GCC^ without affecting the corresponding fl-tRNAs (**Fig. S8a,b**). WB analysis showed that 5’Glu^CTC^-ASO, but not 5’Gly^GCC^-ASO, attenuated glutamate-induced Tau phosphorylation and oligomerization in rat and human neurons (**Fig. 7a**), consistent with reduced oligomeric Tau puncta by immunofluorescence (**Fig. 7b,c**). Similarly, 5’Glu^CTC^-ASO, but not 5’Gly^GCC^-ASO, reduced oligomeric and phosphorylated Tau in Tau P301S_Mut_ human iNs (**Fig. 7d-f**). In Tau-biosensor reporter cells, 5’Glu^CTC^-ASO reversed 5’Glu^CTC^-induced Tau aggregation, as measured by FRET (**Fig. 7g**), with no effect in the absence of 5’Glu^CTC^ overexpression (**Fig. S8c**).

**Fig. 7.**
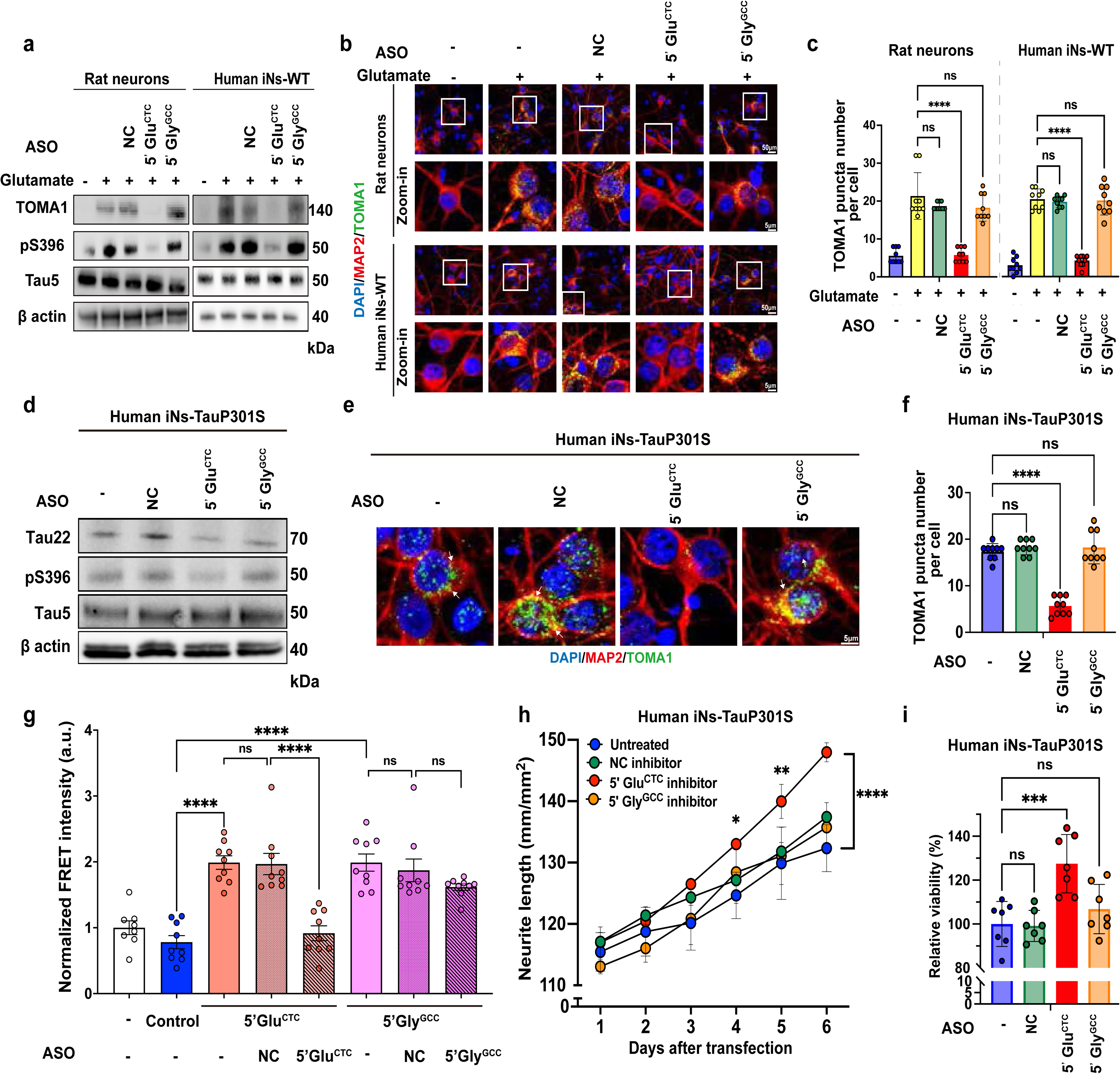
Inhibition of 5’Glu^CTC^ reduces pathological Tau and improves neurite branching and maturation. **a.** Representative WB for indicated Tau species in rat and human neurons transfected with non-template control oligonucleotide (NC), 5’Glu^CTC^ or 5’Gly^GCC^ ASOs, with or without glutamate stress (n = 3/group). **b,c.** Representative IF images (**b**) and quantification (**c**) TOMA1 puncta in rat and human neurons transfected with NC, 5’Glu^CTC^ or 5’Gly^GCC^ ASOs (n = 3/group; ≥250 cells/condition). **d.** Representative WB of indicated Tau species in Tau P301S_Mut_ human neurons transfected with NC, 5’Glu^CTC^ or 5’Gly^GCC^ ASOs (n = 3/group). **e,f.** Representative IF images (**e**) and quantification of TOMA1 puncta in Tau P301S_Mut_ human neurons transfected with NC, 5’Glu^CTC^ or 5’Gly^GCC^ ASOs. **f.** of in (**e**) (n = 3/group; ≥200 cells/condition). **g.** Tau seeding assay in HEK tau-biosensor cells transfected with 5’Glu^CTC^ or 5’Gly^GCC^ plus HMW tau, followed by the ASO treatment 24 hr later. Normalized FRET intensity is shown (n = 9/group). **h,i.** Live-cell imaging of neurite length over 6 days (**h**; n = 3/group) and neuronal viability assessed 6 days after transfection **(i**; n = 7/group) in ASO-treated Tau P301S_Mut_ human neurons. Two-way ANOVA was performed in (**g**). One-way ANOVA with Dunnett’s test was performed in (**c**)(**f**)(**i**). Data are mean ± SEM. ****p < 0.0001, ***p < 0.001, **p < 0.01, *p < 0.05, ns = not significant.

To assess functional effects under subtoxic stress, we monitored neurite length and branching over time. Low-dose glutamate did not impair neurite growth. However, 5’Glu^CTC^-ASO modestly but significantly increased neurite length and branch points (by 2.5-10%) in stressed WT rat and human neurons, with faster responses observed in rat cultures (**Fig. S8d-g**), and similarly enhanced neurite length in Tau P301S_Mut_ human iNs (**Fig. 7h**). The ASO had no effect on non-stressed neurons (**Fig. S8h-j**). Together, reduced Tau oligomerization (**Fig. 7a-d,f,g**) and improved neuronal viability above basal levels in both normal and mutant cultures (**Fig. 7i, Fig. S8k)**, support the potential of 5’Glu^CTC^-targeting strategies for tauopathies.

### Uptake of extracellular 5’tRFs is associated with Tau pathology in recipient neurons

5’Glu^CTC^ is among the most abundant sRNAs in conditioned media (CM) from various cell cultures, including neuronal, and in the cerebrospinal fluid (CSF) and other body fluids^34,77^. Consistent with prior studies^34,78^, fractionation of neuronal CM demonstrated high levels of 5’Glu^CTC^, 5’Gly^GCC^, and 5’Gly^CCC^ in extracellular vesicles (both microvesicles and exosomes), and non-vesicular ribonucleoprotein complexes (RNPs) (**Fig. 8a**). Since extrinsic stress signals and Tau P301S mutation induce tRF accumulation, we examined their effects on extracellular tRFs (ex-tRFs). RNA isolated from CM of rat neurons, naïve or stressed, was used to quantify specific tRFs (**Fig. 8b**). Up to 3-6-fold increases in 5’Glu^CTC^ and 5’Gly^GCC^ were detected in CM from neurons stressed with glutamate, hydrogen peroxide, or 1/2t_max_Aβ (**Fig. 8c**), consistent with intracellular upregulation (**Fig. 2b**). Similarly, CM of Tau P301S_Mut_ human iNs contained elevated 5’Glu^CTC^ and 5’Gly^GCC^ compared to WT iNs (**Fig. 8d**).

**Fig. 8.**
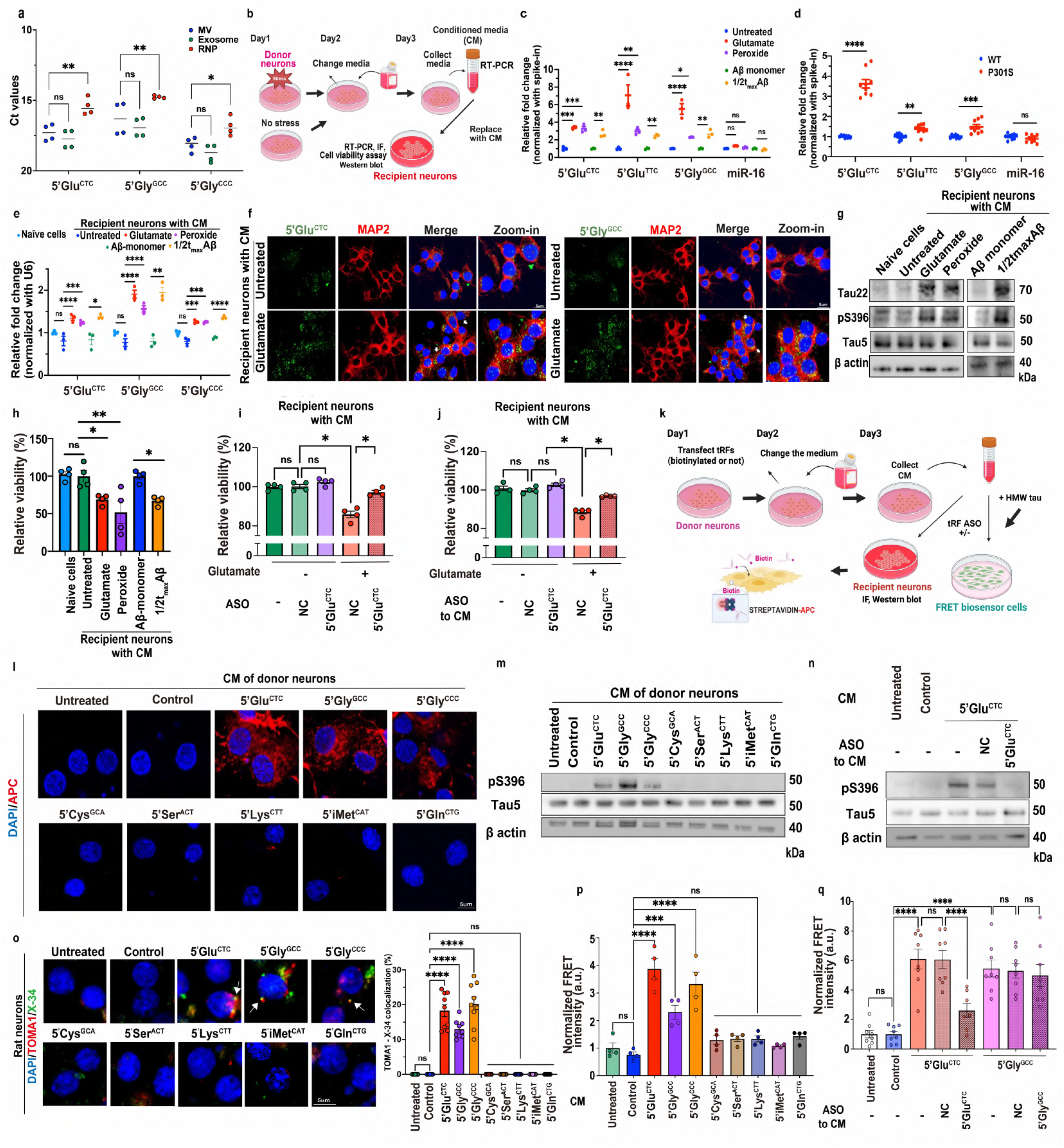
Horizontal transfer of neuronal tRFs is associated with pathological Tau in recipient neurons. **a.** qRT-PCR analysis of extracellular tRFs in microvesicles (MVs), exosomes, and non-versicular ribonucleoproteins (RNPs) from primary neuron conditioned media (CM), shown as Ct values (n = 4/group). **b**. Schematic of CM-transfer experiments in (**c-h**)**. c,d.** qRT-PCR analysis of extracellular-tRFs in CM from stressed rat neurons (**c**) and WT or Tau P301S_Mut_ human neurons (**d**). Normalized to spike-in (n = 3/group). **e,f.** qRT-PCR (**e**) and representative FISH/IF images (**f**) of intracellular 5’tRFs in recipient rat neurons, either naïve or cultured in CM from stressed donor neurons. qRT-PCR normalized to U6 (n = 3/group); 5’Glu^CTC^ or 5’Gly^GCC^ FISH (green), MAP2 IF (red) and nuclei (blue). **g,h.** Representative WB of indicated Tau species (**g**) and viability (**h**) in recipient rat neurons cultured in indicated donor CM (n = 4/group). **i,j.** Viability of recipient neurons cultured in CM from glutamate-stressed donor neurons treated with tRF ASOs or control oligonucleotides before CM collection (**i**), or with ASOs added directly to the CM (**j**) (n = 4/group). **k**. Schematic of 5’tRF-transfer experiments in (**l-q)**. **l.** Representative images of recipient rat neurons, cultured in CM from biotinylated 5’tRF-transfected donor neurons. Recipient cells were stained with allophycocyanin (red) (n = 3/group). **m,n.** Representative WBs of Tau species in recipient neurons cultured with CM from 5’tRF-transfected donor neurons (**m**), or from 5’Glu-overexpressing neurons, with indicated ASOs added to CM (**n**). **o.** Representative TOMA1/X-34 staining and quantification in recipient neurons cultured with CM from 5’tRF-transfected donor neurons. Arrows mark TOMA1/X-34 colocalization. **p,q.** Tau seeding assay in HEK tau-biosensor cells cultured in CM from 5’tRF-overexpressing donor neurons (**p**) or ASOs/control oligonucleotides added to CM (**q**), shown as normalized FRET intensity (p; n = 4/group, q; n = 8/group). One-way ANOVA with Dunnett’s test was performed for **(a), (c), (e), (h-j), (m-n)**. Unpaired two-tailed Student’s *t-test* were performed in **(d).** All graphs show mean ± SEM. ****p < 0.0001, ***p < 0.001, **p < 0.01, *p < 0.05, ns = not significant.

Since ex-sRNA, particularly ex-tRFs, can be taken up by recipient cells, including neurons^27,79,80^, we measured 5’Glu and 5’Gly levels in neurons cultured for 24 hours in CM from naïve or stressed donor neurons (**Fig. 8b**). Recipient cells cultured in CM from stressed neurons showed a 1.3–2-fold increase in 5’Glu^CTC^, 5’Gly^GCC^, and 5’Gly^CCC^ (qRT-PCR, **Fig. 8e**), suggesting interneuronal transfer. Increased FISH signals for 5’Glu^CTC^ and 5’Gly^GCC^ supported this (**Fig. 8f**). In parallel, WB revealed increased oligomeric and phosphorylated Tau in recipient cells, with no change in total Tau (**Fig. 8g**). Recipient neuron viability was reduced compared to cells in fresh media or CM from unstressed neurons (**Fig. 8h**). Transfection of donor neurons with steric-blocker 5’Glu^CTC^-ASO, or neutralization of 5’Glu^CTC^ by adding 5′Glu^CTC^-ASO to the CM, partially rescued the viability of recipient neurons cultured in CM from glutamate-stressed cells (**Fig. 8i,j**), supporting the role of direct tRF transfer.

To test the ex-tRF uptake, we employed biotin-labeled tRFs. Primary neurons were transfected with biotinylated 5’tRFs (**Table S2**), and their CM collected after medium replacement and applied to recipient neurons for 24 hours (**Fig. 8k**). Streptavidin fluorescence demonstrated uptake of 5’Glu^CTC^, 5’Gly^GCC^, and 5’Gly^CCC^ by recipient cultures (**Fig. 8l**). Correspondingly, recipient neurons cultured with CM from 5′Glu^CTC^, 5′Gly^GCC^, or 5′Gly^CCC^ -transfected neurons showed increased pSer396 Tau (**Fig. 8m**), and the effect of 5′Glu^CTC^ was blocked by adding 5′Glu^CTC^-ASO to the CM (**Fig. 8n**). Oligomeric and fibrillar Tau were also detected in recipient cells by TOMA-1 and X-34 staining (**Fig. 8o**).

We next tested whether CM from 5’tRF-overexpressing neurons promotes Tau aggregation in Tau-biosensor cells. CM from 5’Glu^CTC^, 5’Gly^GCC^, and 5’Gly^CCC^-overexpressing neurons, but not other cultures, induced Tau aggregation in recipient cells, quantified by FRET (**Fig. 8p**). Adding titrated 5’Glu^CTC^-ASO to the CM reversed Tau aggregation, whereas 5’Gly^GCC^-ASO had no effect (**Fig. 8q**). Collectively, these findings suggest that specific 5’tRFs, including 5’Glu^CTC^ and 5’Gly^GCC^, generated during persistent neuronal stress, can be horizontally transferred between neurons, promoting accumulation of oligomeric and phosphorylated Tau in recipient neurons.

## Discussion

Previous studies identified sRNAs as the most enriched transcripts in Tau complexes purified from human cells and mouse brains^17,18^. Furthermore, synthetic RNAs of 10–40 nucleotides can induce Tau seeding and promote stable fibril formation *in vitro*^19^. These observations prompted us to systematically investigate endogenous sRNAs altered in AD that may contribute to Tau pathology. Because the sRNome spans diverse lengths and modifications, sequencing biases complicate direct comparisons. To overcome this limitation, we employed hybridization-based sRNA arrays enabling simultaneous detection of thousands of miRNAs, tRNAs, tRFs, and snoRNAs. This analysis identified tRFs are the major dysregulated sRNA class in the AD temporal cortex, with 5’Glu^CTC^ and 5’Gly^GCC^ showing the most consistent upregulation across two independent human brain cohorts, despite limited sample sizes. These findings align with earlier pilot data from other affected brain regions^42^ and re-analyzed AD and FTLD-MAPT^48^ RNAseq datasets. Notably, these tRFs are abundant in both brain tissue and biofluids (CSF, plasma)^77,81^, accumulate with brain aging^82^, and in a mouse neurodegeneration model (**Fig. 3a-f**). Our data indicate that 5’Glu^CTC^ and 5’Gly^GCC^ accumulation may promote Tau pathology through direct binding, increasing hyperphosphorylated, insoluble, and aggregated Tau species.

Remarkably, 5’Glu and 5’Gly also accumulated in rodent and human neurons exposed to diverse stressors, including glutamatergic, oxidative, and Aβ-induced stress, and intrinsic stress associated with the Tau P301S mutation. The robust accumulation of these tRFs across multiple models – *in vitro* and *in vivo*, in rodent and human neurons, and in both wild-type and MAPT-mutant backgrounds – suggests a conserved response to chronic neuronal stress and aging. While tRNA cleavage is a well-established evolutionarily conserved mechanism for translational control during acute stress adaptation^83^, our findings indicate an additional role in chronic neuronal stress and pathology. Consistent with brain aging^82^, we propose that persistent neuronal stress promotes cytoplasmic redistribution of ANG, the primary tRNA-specific ribonuclease, enhancing tRF generation. ANG has been implicated in neurodegenerative disorders, including ALS, PD and AD^84,85^, yet its role in chronic neuronal stress remains poorly understood. 5’Glu^CTC^ and 5’Gly^GCC^ accumulation appears relatively specific to AD, with additional evidence in FTLD-MAPT but not PD, suggesting disease-specific tRF signatures.

Beyond stress-induced ANG-mediated cleavage, accumulation of 5’Glu and 5’Gly may reflect their structural stability, including homo- or heterodimer formation^86^, or protein-binding properties that facilitate sequestration in ribonucleoprotein complexes. Such sequestration may induce structural remodeling, alter activity, or stabilize associated species. Our data demonstrate selective binding of 5’Glu and 5’Gly to Tau within its seeding-competent 295–313 region at the R2/R3 interface, inducing conformational constraint near residue 300. We propose that this promotes Tau misfolding, increases susceptibility to phosphorylation, leading to accelerated oligomerization and fibrillization^73^. 5’Glu and 5’Gly share internal UGGU motifs and dimerization capacity^86^. Although the precise binding determinants, including RNA sequence and higher-order structure, require further investigation, our data support selective Tau binding to 5′Glu and 5′Gly rather than nonspecific polyanionic effects.

The lack of tRF-induced full-length Tau fibrillization *in vitro* is not unexpected, as full-length Tau aggregation is intrinsically slow and faces a higher kinetic barrier *in vitro* than *in vivo*. The jR2R3-P301L fragment was designed to test whether aggregation-inducing tRFs bind specifically to the PHF6-containing R2/R3 segment, a hypothesis supported by BLI-derived affinity measurements. Together, these findings indicate that stress-elevated tRFs promote Tau aggregation via specific binding to a defined Tau interface.

In human iNs and PS19 mouse brains, Tau P301S mutation caused a striking increase in these tRFs, suggesting a potential feedback loop between Tau pathology and tRF dysregulation. Misfolded and aggregated Tau may also serve as a scaffold for tRFs, promoting reciprocal stabilization.

Beyond intracellular functions, tRFs are efficiently secreted and constitute a major class of brain exRNAs. Circulating 5’Glu and 5’Gly are consistently detected in CSF ^50,77,81,87^. Previous studies show tRFs transfer between cells via EVs or direct RNA uptake^27,34,88,35,89^. Our findings suggest that inter-neuronal transfer of 5’Glu and 5’Gly enhances Tau aggregation in recipient cells. Whether ex-tRFs also modulate extracellular Tau species, Tau uptake, or long-range propagation *in vivo* remains unknown. Nevertheless, horizontal inter-neuronal tRF transfer represents an unrecognized mechanism potentially contributing to the spread of Tau pathology.

5’Glu^CTC^ is elevated in aging brain^82^, and our results establish its role in promoting Tau neuropathology. Published estimates^90^, consistent with our data, suggest that abundant tRFs can reach ∼10,000 copies per cell and likely achieve high-nanomolar to low-micromolar concentrations under stress conditions - levels consistent with those inducing Tau fibrillization *in vitro*. Using tRF mimics and ASOs, we demonstrated that modulation of 5’Glu^CTC^ reciprocally affects pathological Tau, neuronal metabolism, and viability. Notably, a 5’Glu^CTC^ steric-blocking ASO reduced pathological Tau in stressed and Tau P301S_Mut_ neurons without affecting fl-tRNA. Beyond Tau regulation, 5’Glu^CTC^ may influence transcription, mRNA stability, translation, and stress granule dynamics^21,36,91,92^. Emerging evidence also links 5’Glu^CTC^ to mitochondrial dysfunction, including disrupted cristae organization, impaired glutaminase-dependent glutamate metabolism, and synaptic deficits, processes highly relevant to AD pathogenesis^82^. These observations suggest that targeting 5’Glu^CTC^ may have therapeutic potential in AD and related tauopathies. However, further studies are needed to optimize specificity, assess off-target effects, and evaluate long-term safety. Although 5′Gly similarly promotes Tau aggregation, its ASO was ineffective, likely reflecting distinct structural or binding properties that may require alternative strategies.

Finally, our findings highlight the need for comprehensive exploration of the tRF landscape in AD and aging. While multiple tRFs are dysregulated in AD brains, this abundant RNA class remains largely understudied. Systematic profiling across brain regions, disease stages, and cell types will be essential. Our sRNA analysis also revealed downregulation of mt-tRFs, predominantly 5’mt-tRFs. A recent PD study similarly identified distinct downregulated mt-tRFs, primarily 3’mt-tRFs, in brain and CSF^50^. Together, these findings suggest a link between mitochondrial dysfunction and tRF dysregulation. Given the central role of mitochondrial impairment in neurodegeneration, mt-tRF biology warrants further investigation.

Previous analyses of Tau aggregates in P301L and WT mice identified enrichment of snRNAs and snoRNAs, with tRNA enrichment also detected despite methods not optimized for tRNA recovery^18^. Our findings suggest that tRFs, particularly 5’Glu and 5’Gly, are important cofactors in Tau pathology. Defining the full spectrum of Tau-sRNA interactions, their structural determinants across cell populations and tauopathies, and *in vivo* causality, will be essential next steps for identifying new therapeutic targets in neurodegenerative disease.

## Resource availability

### Lead contact

Requests for further information and resources should be directed to the lead contact, Dr. Anna M. Krichevsky.

### Materials availability

All unique constructs including 5’tRF mimics and ASOs described in this study can be ordered from IDT (https://www.idtdna.com) using the same sequences.

### Data and code availability

The datasets generated during this study have been deposited at GEO: GSE296695 and GEO: GSE297027 and are publicly available as of the data of the publication. Our original pipeline used for small RNA-seq analysis is available on github (https://doi.org/10.5281/zenodo.18264193).

## Acknowledgements

We thank the NeuroTechnology Studio and Dr. Lai Ding at Brigham and Women’s Hospital for instrument access and data acquisition/analysis support; Drs. Jeremy Koppel and Philippe Marambaud at the Feinstein Institutes for Medical Research for Tau antibodies originally generated by Dr. Peter Davis; Dr. Rachid El Fatimy for Aβ oligomer preparation; Keck Biophysics Facility, a shared resource of the Robert H. Lurie Comprehensive Cancer Center of Northwestern University supported in part by NCI Cancer Center Support Grant #P30 CA060553 for BLI; IMSERC NMR facility at Northwestern University supported by the Soft and Hybrid Nanotechnology Experimental (SHyNE) Resource (NSF ECCS-2025633), NIH 1S10OD012016-01 / 1S10RR019071-01A1, and Northwestern University for NMR. Drs. Erik Uhlmann and Evgeny Deforzh for helpful discussions; the BWH iPSC NeuroHub directed by Dr. Christina Muratore for support with human iN preparation.

This work was supported by the Rainwater Charitable Foundation/ Tau Consortium grants to AMK, SH, and BTH, NIH RFAG090598 to AMK, R01 GM126150 and R01 GM146997 to PI, Cure Alzheimer Fund, the Freedom Together Foundation, and NIH P30AG062421 to BTH, NIH R35GM136411 to SH, and grants from the Ichiro Kanehara Foundation, Uehara Memorial Foundation, and Japan Society for the Promotion of Science to Ami K. Generation of Tau P301S_Mut_ human iPSC lines was supported by the Rainwater Charitable Foundation and Alzheimer’s Disease Research Center grant, NIH P30AG066444 to Dr. Celeste M Karch. Mariana B.M.S. Martins was supported by the Elite Network of Bavaria (Elite Graduate Program Biomedical Neuroscience, S-LW-2016-351/2/58).

## Authors’ contributions

Conceptualization and study design, A.M.K. and Ami K.; investigation, data analysis, and visualization, Ami K.; ThT, TEM, BLI, and NMR, C.T., K.T., and S.H.; bioinformatics and data analysis, L.N. and M.I.B.; tau aggregation assays, assisted by M.B.M. and B.T.H.; 5′tRF pull-down experiments, assisted by P.K.; expression analysis, assisted by A.W., N.K., and Y.A.; FISH and CLIP experiments, assisted by Y.Z.; writing – original draft, Ami K. and A.M.K.; writing – review and editing, all authors.

## STAR★METHODS

## KEY RESOURCES TABLE

| REAGENT or RESOURCE | SOURCE | IDENTIFIER |
| --- | --- | --- |
| **Antibodies** |  |  |
| Mouse monoclonal anti-Tau 5 | Thermo Fisher Scientific | Cat# AHB0042, RRID:AB_2536235 |
| Rabbit polyclonal anti-Tau22 | Millipore | Cat# ABN454, RRID:AB_2888681 |
| Rabbit monoclonal anti-Tau pS396 | Abcam | Cat# ab109390, RRID:AB_10860822 |
| Mouse monoclonal anti-TOMA1 | Millipore | Cat# MABN819, RRID:AB_3068608 |
| Rabbit monoclonal anti-Tau pS202 | Abcam | Cat# ab108387, RRID:AB_10860874 |
| Rabbit polyclonal anti-Tau pS262 | ABclonal | Cat# AP0397, RRID:AB_2771597 |
| Rabbit polyclonal anti-Tau pThr231 | Cell Signaling | Cat# 71429, RRID:AB_2888624 |
| Rabbit monoclonal anti-Tau pThr181 | Cell Signaling | Cat# 12885, RRID:AB_2798053 |
| Rabbit polyclonal anti-MAP2 | Cell Signaling | Cat# 4542, RRID:AB_10693782 |
| Mouse monoclonal anti-PHF-1 | Dr. Peter Davis | N/A |
| Goat Anti-rabbit IgG | Cell Signaling | Cat# 14708, RRID:AB_2798581 |
| Goat Anti-mouse IgG | Cell Signaling | Cat# 14709, RRID:AB_2798582 |
| Goat Anti-rabbit IgG | Cell Signaling | Cat# 7074, RRID:AB_2099233 |
| Goat Anti-mouse IgG | Cell Signaling | Cat# 7076, RRID:AB_330924 |
| Goat Anti-rabbit IgG Alexa Fluor 594 | Invitrogen | Cat# A11012, RRID:AB_2534079 |
| Goat Anti-mouse IgG Alexa Fluor 488 | Invitrogen | Cat# A11029, RRID:AB_2534088 |
| Goat Anti-mouse IgG Alexa Fluor 594 | Invitrogen | Cat# A11032, RRID:AB_2534091 |
| Goat Anti-rabbit IgG Alexa Fluor 488 | Invitrogen | Cat# A11008, RRID:AB_143165 |
| **Biological samples** |  |  |
| Postmortem human temporal cortex (BM36) brain tissues | NIH NeuroBioBank at the Icahn School of Medicine at Mount Sinai | N/A |
| Postmortem human temporal cortex (BM36) brain tissues | NIH NeuroBioBank at Mount Sinai Brain Bank | N/A |
| Postmortem human frontal cortex brain tissues | Massachusetts Alzheimer's Disease Research Center brain bank | N/A |
| **Chemicals, peptides, and recombinant proteins** |  |  |
| Sarkosyl | Sigma-Aldrich | Cat#L5777 |
| NuPage 4–12% Bis-Tris gel | Thermo Fisher Scientific | Cat#NP0322BOX |
| Immun-Blot PDVF membrane | BIO-RAD | Cat#1620177 |
| 0.25% Trypsin-EDTA | Thermo Fisher Scientific | Cat#25200056 |
| 40 µm cell strainer | Sigma-Aldrich | Cat#CLS431750 |
| Neurobasal Medium | Gibco | Cat#21103049 |
| B27 supplement | Gibco | Cat#17504-044 |
| Glutamax | Gibco | Cat#35050061 |
| Poly-D-lysine | Sigma-Aldrich | Cat#P0899 |
| Matrigel | Corning | Cat#354234 |
| Opti-MEM | Thermo Fisher Scientific | Cat#31985070 |
| 1% Lipofectamine 2000 transfection reagent | Thermo Fisher Scientific | Cat#11668019 |
| Trichostatin A | Sigma-Aldrich | Cat#T8552 |
| Sartorius Octet® Streptavidin (SA) Biosensor |  | Cat#18-5019 |
| HiLoad 16/600 Superdex 200 pg column | Cytiva | Cat#28989335 |
| Dynabeads™ MyOne™ Streptavidin C1 | Thermo Fisher Scientific | Cat#65001 |
| Formvar-coated carbon film | Electron Microscopy Science | Cat#FCF-200-Cu |
| RNase A | Thermo Fisher Scientific | Cat#12091021 |
| RNase T1 | Thermo Fisher Scientific | Cat#AM2283 |
| N-PER Neuronal Protein Extraction Reagent | Thermo Fisher Scientific | Cat#87792 |
| Complete, Mini, EDTA-free Protease Inhibitor Cocktail | Millipore | Cat#11836170001 |
| NeuroMag transfection reagent | OZ Biosciences | Cat#NM51000 |
| Anti-digoxigenin-alkaline phosphatase | Roche | Cat#11207733910 |
| CDP-Star ready-to-use | Roche | Cat#12041677001 |
| X-34 | Cell Signaling | Cat#74193 |
| Phenol:chloroform:isoamyl alcohol (25:24:1) solution | Thermo Fisher Scientific | Cat#J62336-AN |
| Propidium iodide | Sigma-Aldrich | Cat#P4170 |
| RIPA buffer | Boston Bio-Products | Cat#BP-115X |
| ROCKi | StemCell Technologies | Cat#Y-27632 |
| VECTASHIELD HardSet Antifade Mounting Medium with DAPI | Vector | Cat#H-1500-10 |
| 1X protease and phosphatase inhibitors | Cell Signaling | Cat#5872 |
| **Critical commercial assays** |  |  |
| AlamarBlue HS Cell viability | Thermo Fisher Scientific | Cat# A50100, |
| Pierce™ BCA Protein Assay Kits | Thermo Fisher Scientific | Cat# 23225 |
| Vybrant™ DyeCycle™ Violet Stain | Thermo Fisher Scientific | Cat# V35003 |
| ECL chemiluminescence reagents | Genesee Scientific | Cat#20-300 |
| ViewRNA cell Plus Assay Kit | Invitrogen | Cat#88-1900-99 |
| miRCURY LNA SYBR Green PCR kit | QIAGEN | Cat#339345 |
| miRCURY LNA RT kit | QIAGEN | Cat#339340 |
| **Deposited data** |  |  |
| Microarray datasets generated during this study | This paper | GSE296695, GSE297027 |
| Our original pipeline used for small RNA-seq analysis | This paper | https://doi.org/10.5281/zenodo.18264193 |
| **Experimental models: Cell lines** |  |  |
| *MAPT*<sup>P301S</sup> human iPSC line | Karch CM et al<sup>93</sup> | Donor ID: F0510 |
| *MAPT*<sup>WT</sup> human iPSC line | Karch CM et al<sup>93</sup> |  |
| Rat primary cortical neurons from E18 SAS Sprague Dawley pups |  | Charles River and Taconic |
| **Experimental models: Organisms/strains** |  |  |
| Mouse: B6;C3- Tg(Prnp-MAPT*P301S)PS19Vle/J | The Jackson Laboratory | Jax strain # 008169 |
| **Software and algorithms** |  |  |
| ImageJ | Schindelin et al | https://imagej.net/ij/ |
| Prism 10 | GraphPad | https://www.graphpad.com/ |
| R (version 4.1.3) | GNU General Public License | https://www.R-project.org/ |
| LIMMA (version 3.50.3) | Bioconductor | http://bioinf.wehi.edu.au/limma |
| BioRender | BioRender | https://www.biorender.com/ |

## EXPERIMENTAL MODEL AND STUDY PARTICIPANT DETAILS

### Brain tissues

Frozen human post-mortem brain specimens were obtained from the Harvard Brain Tissue Resource Center, NIH NeuroBioBank at the Icahn School of Medicine at Mount Sinai and the Massachusetts Alzheimer’s Disease Research Center brain bank. The specimens were used in accordance with the policies of Brigham and Women’s Hospital’s institutional review board (#2011B000278/BWH). Detailed neuropathological and demographic information about the patient samples used is included in **Supplementary Table S1**. Mouse brain specimens were prepared from WT and PS19 mouse (B6;C3-Tg(Prnp-MAPT*P301S)PS19Vle/J). Mice were maintained on a 12:12-h light/dark cycle (7:00 am on/7:00 pm off) with food and water provided ad libitum before experimental procedures. The research complies with all relevant ethical regulations. Experiments involving animals were carried out in accordance with the recommendations in the U.S. National Institutes of Health Guide for the Care and Use of Laboratory Animals. The protocols (2016N000113, 2016N000114) were approved by the Institutional Animal Care and Use Committee at Brigham and Women’s Hospital.

### Rat primary cortical neuron culture

Rat primary cortical neuron cultures were prepared from E18 SAS Sprague Dawley pups (Charles River and Taconic). Brain tissues were dissected, dissociated enzymatically by 0.25% Trypsin-EDTA (Thermo Fisher Scientific), triturated with fire-polished glass Pasteur pipettes, and passed through a 40 μm cell strainer (Sigma-Aldrich, Cat# CLS431750). After counting, neurons were seeded onto poly-D-lysine (Sigma-Aldrich) coated cell culture plates at 80,000 cells/cm2 in Neurobasal medium (Gibco, Cat#21103049) supplemented with 1X B27 (Gibco, Cat#17504-044) and 0.25X GlutaMax (Gibco, Cat#35050061). Half medium was changed every 4 days until use.

### Human iPSC-derived neurons

24-well plates, 96-well plates, and 10cm dishes (Corning) were coated with Matrigel (Corning, Cat#354234) solution (0.2 mg/mL in DMEM/F12) and poly-o/laminin for 1.5 h at 37 °C. Then, the Matrigel solution was completely removed, and PBS (Gibco) was added. The plates and dishes were incubated at 37°C shortly before use. Frozen day 4 iPSC-derived neurons were thawed in pre-warmed resuspension media per vial, which was composed of NBM (Neurobasal medium, 0.5x MEM-NEAA, 1x GlutaMAX, 0.3% dextrose), 1:50 B27, 1:10000 DOX, puromycin, 1:1000 BDNF/CNTF/GDNF and 1:1000 ROCKi (StemCell Technologies) and were kept in a warm metal bath to facilitate the thawing. After gently mixing by inverting the tube, viable cell concentration was determined using trypan blue (Bio-Rad). The cells were centrifuged (250 g, 5 min, room temperature), resuspended in day 4 medium, and plated at the density of 5 × 10^4^ cells/cm^2^. On days 7/10/14/18, two thirds of the conditioned medium was replaced with pre-warmed medium (NBM, 1:50 B27, 1:10000 DOX, puromycin, 1:1000 BDNF/CNTF/GDNF). The following iPSC lines were used: lines derived from *MAPT P301L* donor (donor ID: F0510) which were either mutated to *MAPT P301S* using CRISPR/CAS9 (F0510.2Δ3B5) or corrected to MAPT WT (F0510.2Δ3B3, 4, E10), provided by Dr. Celeste M Karch^93^. The iPSC-derived neurons were generated following review and approval through Brigham and Women’s Hospital Institutional Review Board (#IBR2015P001676).

## METHOD DETAILS

### RNA isolation and quantitative RT-PCR (qRT-PCR)

Total RNA was extracted from cells and tissues using Trizol (Invitrogen), and from CM and its fractions using miRCURY protocol, with on-column DNase treatment (Qiagen, Germany). RNA quality was assessed using Thermo Scientific Nanodrop 1000 Spectrophotometer (RRID:SCR_016517). For tRF analysis, 200ng of the total RNA was reverse transcribed using miRCURY LNA RT kit (QIAGEN, Cat#339340). The resulting cDNA libraries were diluted 80 times and used for the analysis of gene expression by qPCR using miRCURY LNA SYBR Green PCR kit (QIAGEN, Cat#339345) with miR-CURY LNA^TM^ primers (YP00203907 for U6; YP02114672 for snoRD115, YP00206035 for miR-132-3p; YP00204170 for miR-212-3p; YP00204521 for miR-99a-5p) (QIAGEN) (**Supplementary Table S4**). qPCR was performed on Applied Biosystems ViiA 7 Real-Time PCR System (RRID:SCR_023358). The cycling conditions were 95 °C for 10 min, 50 cycles of 95 °C for 15 s, and 60 °C for 1 min following dissociation analysis. The fold changes in gene expression were calculated by the ΔΔCt method. The expression of sRNA was normalized to miR-99a and U6 snRNA. As our target subset of 5’ half of tRNAs are minimally modified (and so-called “hard-stop” modifications, including 1-methyladenosine (m^1^A), 1-methylguanosine (m^1^G), 2,2,-dimethylguanosine (m^2,2^G), and 3-methylcytidine (m^3^C), are primarily found in 3’- and not 5’-tRFs and that most of the modifications are found after position 31 where cleavage occurs), our qRT-PCR method largely bypasses the barriers for reverse transcription.

### Fractionation of cells and conditioned media (CM)

Sarkosyl-soluble and insoluble Tau fractions were prepared based on a published protocol^94,95^. Fractionation of CM was performed based on a published protocol^34^. Approximately 100 ml CM was used as input for exRNA isolation. CM was centrifuged at 300×g, 4 °C for 10 min, following by the additional centrifugation at 2000×g, 4 °C for 15 min, to remove cells and cell debris. 5 μl of SUPERase In RNase Inhibitor (Ambion) was added to the supernatants per 10 ml media. The media was then filtered sequentially through the 2 μm filter (GE Healthcare, UK), 0.8 μm filter (EMD Millipore, MA), and 0.22 μm filter (EMD Millipore), with no/minimal pressure applied. The filtrate was split to 15 ml per sample and further filtered through the 0.02 μm filter (GE Healthcare) with up to 75 psi pressure applied. The fractions collected on 0.02 μm filters were lysed with 900 μl lysis solution. The last flow-through fractions of 0.02 μm filters were pooled together (up to 30 ml) and concentrated ∼60 times using 3 kDa Amicon Ultra Centrifugal Filters (EMD Millipore) at 4000×g, 4 °C, for 60 min.

### Northern blot

Total RNA was separated on 15% TBE-urea gels (ThermoFisher Scientific) and transferred onto positively charged nylon membranes (Sigma 11417240001). The membranes were cross-linked by UV irradiation and hybridized overnight at 40°C with digoxigenin (DIG)-labeled DNA probes detecting specific tRFs, in DIG Easy Hyb solution (Roche). After low stringency washes (twice with 2× SSC/0.1% SDS at room temperature) and high stringency wash (once with 1× SSC/0.1% SDS at 40°C), the membranes were blocked in blocking reagent (Roche) for 30 min at room temperature, probed with anti-digoxigenin-alkaline phosphatase (Roche) for 30 min, and washed with 3x TBS-T. Signals were developed using CDP-Star ready-to-use (Roche) and detected using the BioRad ChemiDoc Touch Imaging System (RRID:SCR_021693) according to the manufacturer’s instructions. DIG-labeled oligonucleotide probes were synthesized by Integrated DNA Technologies (IDT). The sequences of the probes are listed in **Supplementary Table S5**. Densitometry analyses were performed using ImageJ (RRID:SCR_003070).

### Western blot

Cells were lysed in RIPA buffer (Boston Bio-Products) supplemented with Complete, Mini, EDTA-free Protease Inhibitor Cocktail (Millipore Sigma). Protein concentrations were quantified using the MicroBCA Protein assay kit (ThermoFisher Scientific). Equal amounts of total proteins (30μg per sample) were separated by SDS PAGE in 4-12% Bolt Bis-Tris Plus gels (ThermoFischer Scientific) and transferred to 0.2 μm Immun-Blot PDVF membrane (BIO-RAD). After blocking with 5% (w/v) fat free milk in tris-buffered saline with 0.1% Tween (TBS-T, Boston Bioproduct) for 1h, membranes were incubated with primary antibodies: anti-Tau 5 (Thermo Fisher Scientific Cat# AHB0042), 1:5000 dilution; anti-Tau22 (Millipore Cat# ABN454), 1:1000 dilution, anti-Tau pS396 (Abcam Cat# ab109390), 1:5000 dilution; anti-Tau pS202 (Abcam Cat# ab108387), 1:5000 dilution; anti-Tau pS262 (ABclonal Cat# AP0397), 1:500 dilution; anti-Tau pThr231 (Cell Signaling Cat# 71429, RRID:AB_2888624), 1:1000 dilution; anti-Tau pThr181 (Cell Signaling Cat# 12885), 1:1000 dilution; anti-TOMA1 (Millipore Cat# MABN819), 1:100 dilution; anti-PHF-1, originally created by Dr. Peter Davis and kindly provided by Dr. Jeremy Koppel and Dr. Philippe Marambaud at the Feinstein Institutes for Medical Research, 1:100 dilution) overnight. Blots were washed and incubated with secondary antibodies (1:5000 dilution) for 2 h at room temperature. After washing, bands were visualized with ECL chemiluminescence reagents (Genesee Scientific) using the ChemiDoc Imaging System (Bio-Rad). Densitometry analyses were performed using ImageJ (RRID:SCR_003070).

(Sigma-ldrich), Nicotinamide mononucleotide (Sigma-Aldrich), PDD

### Immunofluorescence staining

Cells were seeded onto poly-D-lysine (Sigma-Aldrich) coated coverslips and briefly washed with PBS 3 times, fixed with 4% paraformaldehyde (PFA), permeabilized with 0.5% Triton X-100 in PBS, and blocked in 5% BSA for 1h. The cells were then incubated with primary antibodies overnight at 4°C and further incubated with secondary antibodies (goat anti-rabbit Alexa Fluor 488, Cat# A11008; goat anti-rabbit Alexa Fluor 594, Cat# A11012,, goat anti-mouse Alexa Fluor 488, Cat# A11029; goat anti-mouse Alexa Fluor 594, Cat# A11032; Invitrogen, 1:500 dilution). The nuclei were stained with DAPI (1:1000). The staining was visualized and imaged using a Zeiss LSM 880 with Airyscan Confocal Laser Scanning Microscope (RRID:SCR_020925).

### X-34 staining

After fixation with 4% PFA, cells were incubated in 25 µM X-34 staining solution for 30 min at room temperature in the dark. Cells were then rinsed three times with ddH₂O and incubated in 50 mM NaOH/ 80% ethanol buffer for 2 min, followed by three additional washes with ddH₂O. Immunostaining was subsequently performed. Fluorescence imaging was acquired using a Zeiss LSM 880 Airyscan confocal laser scanning microscope (RRID:SCR_020925).

### Fluorescence *in situ* hybridization (FISH) and immunofluorescence for cell cultures

RNA FISH was performed using the ViewRNA cell Plus Assay Kit (Invitrogen, Cat#88-1900-99) following the manufacturer’s instructions. Alexa Flour 488 (green), 594 (red), and 647 (magenta)-conjugated probes were custom-made by ThermoFisher. 4 × 10^4^ cells per well were cultured on poly-D-lysine (Sigma-Aldrich, Cat#P0899) coated coverslips in a 24-well plate (Corning), fixed with 4% PFA, and permeabilized with 0.5% Triton X-100 in PBS. For the FISH-Immunofluorescence staining combination, the cells were blocked in 5% BSA for 1h and go through overnight incubation at 4°C with primary antibody; anti-TOMA1 (Millipore Cat#MABN819, RRID:AB_3068608), 1:100 dilution; anti-MAP2 (Cell Signaling Technology Cat#4542, RRID:AB_10693782), 1:1000 dilution before hybridization with FISH probes. After three washes with RNase inhibitor-containing PBS, the cells were hybridized with probes specific for 5’Glu^CTC^ and 5’Gly^GCC^ for 2 h at 40 °C. The sequences of the probes are provided in **Supplementary Table S5**. For background assessment, the probes were omitted in parallel experiments. The cells were further washed with Cell Plus RNA Wash buffer; the tissue slides were washed with 2X SSC for 5 times at 40°C. The cells were hybridized sequentially with PreAmplifer, Amplifer, and Label probe solution for 1 h at 40 °C. For the FISH-Immunofluorescence staining combination, the slides were incubated with secondary antibodies (goat anti-mouse Alexa fluor 488, Cat#A11029, RRID:AB_2534088; goat anti-rabbit Alexa fluor 488, Cat#A11008, RRID:AB_143165; goat anti-mouse Alexa fluor 594, Cat#A11032, RRID:AB_2534091; goat anti-rabbit Alexa fluor 594, Cat#A11012, RRID:AB_2534079; Invitrogen, 1:5000 dilution) for 2h at room temperature. They were further washed with PBS three times, stained with 1 × DAPI (4’,6-diamidino-2-phenylindole), mounted in a mounting media (Vector, #H-1500-10), and observed under Zeiss LSM 880 with Airyscan Confocal Laser Scanning Microscope (RRID:SCR_020925). Quantification of the colocalization was determined by ImageJ (RRID:SCR_003070).

### Fluorescence *in situ* hybridization (FISH) and immunofluorescence for brain sections

For the mouse hippocampus analysis, mice were sacrificed by CO_2_ exposure, followed by cervical dislocation. Brains were extracted and cryoprotected in 30% sucrose for 4-6 h RT, embedded in optimal cutting temperature compound (OCT), sectioned at 15μm thickness, and fixed in acetone for 5 min at − 20 °C. After three washes with RNase inhibitor-containing PBS, the tissue slides were permeabilized with 0.3% Triton-X 100 in PBS for 5 min on ice. The slides were fixed in 4% PFA, treated with 0.3% Triton-X 100 solution, and blocked with 5% BSA. Tissue slides were placed into acetone and then treated with antigen repair solution (20 mg/ml Proteinase K, 1 M Tris-HCl (pH=8.0), 0.5M EDTA (pH=8.0)) for 5min at 37°C. For the FISH-Immunofluorescence staining combination, the slides the step of overnight incubation at 4°C with primary antibody: anti-TOMA1 (Millipore Cat#MABN819, RRID:AB_3068608), 1:100 dilution; anti-MAP2 (Cell Signaling Technology Cat#4542, RRID:AB_10693782), 1:1000 dilution) was included before hybridization with FISH probes. The slides were hybridized with probes specific for 5’Glu^CTC^ and 5’Gly^GCC^ for 2 h at 40 °C, and washed with 2X SSC for 5 times at 40°C. The slides were hybridized sequentially with PreAmplifer, Amplifer, and Label probe solution for 1 h at 40 °C. For the FISH-Immunofluorescence staining combination, the slides were incubated with secondary antibodies (goat anti-mouse Alexa fluoro 488, goat anti-rabbit Alexa fluoro 488, goat anti-mouse Alexa fluoro 594, goat anti-rabbit Alexa fluoro 594, Invitrogen, 1:5000 dilution) for 2h at room temperature. They were further washed with PBS three times, stained with 1 × DAPI (4’,6-diamidino-2-phenylindole), mounted in a mounting media (Vector Cat#H-1500-10), and observed under Zeiss LSM 880 Confocal Laser Scanning confocal microscope. Quantification of the colocalization was determined by Image J pro 6.0.

### RNA-protein crosslinking and immunoprecipitation (CLIP)

CLIP was performed based on previous published protocol^96^, with the following modifications. To covalently crosslink proteins to nucleic acids, 2 × 10^7^ cells or homogenized mouse hippocampi were exposed to UV irradiation (200mJ/cm^2^) for 2 minutes. The cells and tissues were then lysed in 100 ul RNA-binding protein immunoprecipitation (RIP) lysis buffer supplemented with 0.5 ul Protease Inhibitor Cocktail (Millipore Sigma) and RNase Inhibitor (0.002 U/ml). The lysates were incubated on ice for 5 minutes. In parallel, immunoglobin-coated protein magnetic beads (Thermo Fisher Scientific) were washed 2 times with RIP buffer, mixed with antibodies: anti-Tau 5 (Thermo Fisher Scientific Cat# AHB0042, RRID:AB_2536235); anti-Tau pS396 (Abcam Cat# ab109390, RRID:AB_10860822); anti-PHF-1, originally created by Dr. Peter Davis and kindly provided by Dr. Jeremy Koppel and Dr. Philippe Marambaud at the Feinstein Institutes for Medical Research, 1:100 dilution) and incubated for 30 minutes at room temperature with rotation. Then, RIP lysates were centrifuged at 14,000 rpm for 10 minutes at 4°C. After supernatant was removed, beads-antibody complexes were added and incubated for overnight at 4°C. Following incubation, the bead-lysate mixture was washed 15 times intensively with cold RIP buffer, and the samples were treated with DNase I treatment (Promega) for 30 minutes at 37°C, followed by proteinase K (10% SDS and 10mg/ ml proteinase K in RIP buffer) treatment for 30 minutes at 55°C with shaking to digest the protein. The beads were collected with a magnetic stand, and the supernatants were transferred to a new tube. RNA was further isolated using 400 ul phenol:chloroform:isoamyl alcohol (25:24:1) solution (Thermo Fisher Scientific, Cat#J62336-AN), precipitated with isopropanol, and resuspended in RNase-free water.

### 5’tRF mimics, antisense oligonucleotides (ASOs) and cell transfection

Transfections of DIV10 rat primary neurons and DIV21 human iNs^43^ were performed using NeuroMag transfection reagent (OZ Biosciences, Cat#NM51000). Synthetic 5’tRFs (synthesized, desalted, and high-performance liquid chromatography (HPLC)-purified by IDT, resuspended in nuclease-free water to generate stock solutions and stored at -80°C until use. Sequences in Table S2) were transfected at a final concentration of 20 nM to rat neurons and 2 nM final concentration to human iNs. 5’tRFs ASOs (2’OMe-ZEN-containing, IDT, Sequences in Table S3) were transfected at 2-8 nM final concentration for rat neurons and 0.2-0.8 nM final concentration for human iNs. Corresponding control oligonucleotides of the same chemistries were carried out in parallel. Transfection efficiency of these oligonucleotides was 90–95%. For tRF neutralization in CM, the CM were collected, cleared by filtration using 0.22 μm filter (EMD Millipore), and the ASOs added at a concentration of 50 nM.

### Pull down of 5’tRF-bound proteins

Rat primary neurons transfected with biotinylated 5’RNAs were subjected to UV irradiation (200mJ/cm2) to covalently crosslink proteins to nucleic acids. Cells were then lysed with N-PER Neuronal Protein Extraction Reagent (Thermofisher scientific, Cat#87792) supplemented with Protease Inhibitor Cocktail (Millipore Sigma), phosphatase inhibitor, and RNase Inhibitor (Promega). Protein concentrations were measured using Pierce BCA Protein Assay Kit (Thermo Scientific, Cat# 23225) according to the manufacture’s protocol. Equal amounts (∼5 mg) of lysates were used for each pull-down. For each pull down, 60 µL of streptavidin-conjugated magnetic beads (Dynabeads™ MyOne™ Streptavidin C1, ThermoFisher, Cat# 65001) were washed twice with 1 mL of wash buffer 1 (10 mM Tris.HCl pH 7.4, 100 mM NaCl, 1mM EDTA, pH 8.0, 0.1% NP40). Washed beads were then mixed with 300 µL of the lysates and 300 µL of wash buffer 1 supplemented with protease inhibitor, phosphatase inhibitor, and RNase inhibitor. The mixture was rotated overnight at 4°C.

Following incubation, the bead-lysate mixture was washed 5 times with wash buffer 1, twice with wash buffer 2 (10 mM Tris.HCl pH 7.4, 300 mM NaCl, 1mM EDTA, pH 8.0, 0.1% NP40), and once with wash buffer 1. All the liquid was carefully removed, and the samples were treated with a mixture of RNase A (500 U/mL, ThermoFisher, Cat#12091021) and RNase T1 (200 U/mL, ThermoFisher, Cat#AM2283) for 30 min at 37 C. Proteins bound to biotinylated 5’tRFs were eluted in 2X SDS gel loading buffer. The beads were collected with a magnetic stand, and the supernatants were used for Western blot detection of Tau variants with primary antibodies of anti-Tau pS396 (Abcam Cat# ab109390, RRID:AB_10860822) and anti-Tau 5 (Thermo Fisher Scientific Cat# AHB0042, RRID:AB_2536235).

### Small RNA labeling, microarray hybridization, and analysis

Total RNA was dephosphorylated to remove residual phosphate and cyclic phosphate groups from the 3′ ends, exposing the 3-OH termini. The reaction was terminated by heating the samples at 70°C for 3 min, followed by rapid cooling on ice. The samples were then denatured at 100°C and RNA end-labeling was performed using T4 RNA ligase in the presence of 50 mM pCp-Cy3/Cy5, ligase buffer, and BSA, with incubation overnight at 16 °C. Subsequently, the labeled RNAs were hybridized to human/mouse sRNA microarrays (GPL31220/GPL31221, Arraystar). Slides were scanned on Agilent G2505C microarray scanner (Agilent, California, U.S.A.).

Microarray data were processed using Agilent Feature Extraction Software (RRID:SCR_014963) (version 11.0.1.1). Raw signal intensities log-2 transformed, and quantile normalized using the Agilent Technologies GeneSpring GX (RRID:SCR_010972) v12.1 software package (version 11.0.1.1). After normalization and quality control, the probe sets flagged as either “Present” (P) or “Marginal” (M) were retained. For genes represented by multiple probes, expression values were summed to yield a single expression value for all tRF isocoders. Differentially expressed tRFs were identified using the LIMMA (RRID:SCR_010943) (version 3.50.3) in R (version 4.1.3), with a |fold change (FC)| threshold > 1.5 and p-value < 0.05 considered statistically significant. Finally, hierarchical clustering, volcano plots, and scatter plots were generated to visualize distinct tRF expression patterns across samples. Microarry datasets were deposited into the Gene Expression Omnibus (GEO) Repository with accession numbers GSE296695 and GSE297027.

### Reanalysis of published small-RNA-Seq data

Publicly available small RNA sequencing datasets from frontal and prefrontal cortex were analyzed, including Alzheimer’s disease samples (controls/early-stage, n = 6; late-stage, n = 6) (GSE48552), frontotemporal dementia with MAPT mutations (FTD-MAPT; n = 13) with controls (n = 12) (E-MTAB-12731, batch 1) and Parkinson’s disease (PD; n = 29) and controls (n = 36) (GSE72962, GSE64977). Raw sequencing data were obtained from the Sequence Read Archive and ArrayExpress and trimmed to remove adapter sequences according to each library construction protocol. Following adapter trimming and removal of contaminant sequences (UniVec), reads were sequentially aligned to ribosomal RNA (rRNA), transfer RNA (tRNA) with subsequent tRNA fragment (tRF) classification, microRNA/snoRNA/scaRNA, piRNA, and finally protein-coding mRNA, long noncoding RNA (lncRNA), and miRNA hairpin references. Reference annotations were obtained from RNAcentral (rRNA), GtRNAdb (tRNA), miRBase (miRNAs), piRBase (piRNAs), and Ensembl (snRNA, snoRNA, other ncRNAs, mRNA, and lncRNA). Using this multi-step alignment workflow, an average of 90–95% of reads were successfully mapped across samples. The complete alignment pipeline is available on GitHub (https://doi.org/10.5281/zenodo.18264193). tRNA fragments were classified following the tRAX framework (http://trna.ucsc.edu/tRAX/, https://github.com/UCSC-LoweLab/tRAX)^97,98^ based on read position relative to mature tRNA boundaries: reads mapping within 10 nucleotides of the 5′ end were classified as 5′ tRFs, those within 10 nucleotides of the 3′ end as 3′ tRFs, and reads mapping to internal regions that did not overlap either end were classified as internal tRFs. Reads spanning both 5′ and 3′ boundaries were classified as full-length tRNAs. However, such reads were rare given the short-read length (<50 nt) of small RNA-seq libraries. Following alignment, low-abundance RNA species (<∼10 reads) were filtered and counts for individual tRF isodecoders were summed prior to analysis. Differential expression analysis was performed using DESeq2^99^ with normalization based on total mapped reads. Statistical significance was defined as a false discovery rate (FDR) < 0.05 with a fold change >1.5 or <0.5 (|log2 fold change| > 0.585). Figures were generated in R using ggplot2.

### Preparation of oligomeric high molecular weight tau (HMW Tau) seeds from human brain

Frozen human AD tissue (5g) was thawed on wet ice, gray matter was dissected and immediately homogenized in PBS containing 1X protease and phosphatase inhibitors (Cell Signaling, Cat#5872) + 2 µM Trichostatin A (Sigma, Cat#T8552) in a 15-ml glass Dounce homogenizer. The homogenate was centrifuged at 10,000g for 10 min at 4 °C. The supernatants were collected, aliquoted and stored at − 80 °C until further processing. The total volumes of soluble brain extracts were separated by size exclusion chromatography (SEC) as previously described^100^ on a single HiLoad 16/600 Superdex 200 pg column (Cytiva, Cat#28989335) in PBS (Sigma-Aldrich, Cat#P3813, filtered through a 0.2-um membrane filter), at a flow rate of 0.5 ml/min using an AKTA purifier 10 (GE Healthcare). For each run, 5 ml of soluble brain extract was loaded onto the column, and 56 fractions of 2 ml were collected. Fractions of 2 ml were retrieved, analyzed by western blots, and their bioactivity was assessed with Tau seeding assays. The fractions containing seeding-competent, oligomeric high-molecular-weight Tau species which refers to high molecular weight (HMW) tau previously described as Fractions 2-3-4 in Dujardin et al.^100^ were stored at − 80 °C until further use for *in vitro* seeding assay.

### Cell based tau-seeding assay

For tau seeding assessment, FRET-biosensor HEK293 cells stably expressing the PS19-mutant tau repeat domain conjugated to either cyan fluorescent protein (CFP) or yellow fluorescent protein (YFP) (TauRD-P301S-CFP/YFP) were employed^100,101^. Briefly, cells were plated on 96-well plates (Costar, previously coated with 1:20 poly-d-lysine) at a density of 20,000 cells per well and cultured for 24 h at 37 °C, 5% CO_2_ in Dulbecco’s modified Eagle medium (DMEM), 10% fetal bovine serum (FBS) and 1% penicillin–streptomycin. HMW tau extracts (8 ng total tau per well) were incubated with 1% Lipofectamine 2000 transfection reagent (Thermo Fisher, Cat#11668019) in Opti-MEM (Thermo Fisher, Cat#31985070, final volume of 50 µl per well) for 20 min at RT before being added to the cells. After 24 h, cells were collected for subsequent flow cytometer analysis of seeding, i.e., FRET signal. The medium was removed, 50 µl trypsin 1X was added to each well for 5 min at 37 °C, and the reaction was stopped by adding 150 µl fresh medium. Cells were transferred to 96-well U-bottom plates (Corning), pelleted at 1500 rpm for 10 min, resuspended in 2% PFA in PBS for 20 min at RT and pelleted again at 1500 rpm for 10 min. Cells were finally resuspended in 150 µl per well PBS and run and analyzed on the MACSQuant VYB (Miltenyi) flow cytometer as previously described. For each well, tau seeding value was calculated by multiplying the percentage of FRET-positive cells by the median fluorescence intensity of that FRET-positive population (integrated FRET density or IFD). Each sample was loaded in triplicate and three experiments were performed independently.

### Preparation of Tau construct

The Tau construct 0N4R-P301L (residues 1–441, 383 amino acids in length, with residue 44 followed by residue 103 based on the 2N4R numbering scheme), which lacks both N-terminal inserts encoded by exons 2 and 3 and harbors the missense mutation P301L, was expressed recombinantly in *E. coli* and purified following previously reported methods^72^. The truncated tau construct jR2R3-P301L (residues 295–313: DNIKHVPGGGSVQIVYKPV), spanning the R2/R3 domains of human full-length tau, was also used for *in vitro* experiments (Thioflavin T assay, TEM, NMR, and BLI) in this work to assess the effects of tRF on Tau aggregation. The fibrils formed by this construct adopt a U-shaped fold found within the strand-loop-strand structure formed by this segment in fibrils in 4R tauopathies and recruits naïve 4R Tau for templated aggregation^72,74^, making it an optimal construct for evaluating the efficacy of different co-factors in promoting Tau pathology. Non-isotope-enriched peptides were purchased from GenScript, while the ^15^N-enriched peptides for NMR studies were synthesized by the Peptide Synthesis Core at Northwestern University using solid-phase peptide synthesis (SPPS), with isotope labeling on valine, leucine, and isoleucine. The lyophilized powders were dissolved to 1 mM stock solutions using 20 mM ammonium acetate, 50 mM NaCl, pH 7.4 (calibrated with HCl and NaOH) buffer and stored at −80° C or used immediately.

### Thioflavin T aggregation assay *in vitro*

All experiments were performed with a Tecan Infinity Nano+ plate reader. In each well 50 uM jR2R3-P301L tau peptide, 20 µM ThT, and 12.5 µM of the co-factor (heparin or tRNA fragments) in buffer were distributed into a 384-well plate (Corning low volume non-binding surface black with clear flat bottom) to a total volume of 20 µL. The plate reader temperature was set to 37 °C and allowed to equilibrate, the samples were orbital shaken under the programmed shaking speed 3 in between measurements. ThT fluorescence intensity was measured at (excitation=430 nm, emission=485 nm) every 5 minutes until a plateau was reached. These experiments were done in triplicate at least three times with independent samples.

### Transmission electron microscopy (TEM) analysis

For transmission electron microscopy (TEM) analysis, 5 μL of fibril samples were applied to a glow-discharged 200 mesh copper grid with a Formvar-coated carbon film (Electron Microscopy Science, Cat#FCF-200-Cu) for 1 min and blotted dry with filter paper. Samples were stained with 5 μL 1.5 w/v % uranyl acetate and immediately blotted dry. An additional 5 µL of uranyl acetate was added for 1 min and blotted dry. Samples were analyzed using JOEL 1400 TEM in the Northwestern University Atomic and Nanoscale Characterization Experimental Center (NUANCE) operated at 120 kV and room temperature. The fibrils were then imaged with a Gatan OneView^TM^ camera at the indicated magnifications.

### NMR spectroscopy

1H/15N SOFAST-HMQC experiments were recorded using a Bruker Neo 600 MHz system with a QCI-F cryoprobe. Experiments used 200 uM 15N-labeled jR2R3-P301L tau peptides mixed with tRF co-factor (4:1 molar ratio) in 20 mM ammonium acetate, 50 mM NaCl, pH 7.4, supplemented with 10% D2O. Spectra were recorded at 298 K, with a recycle delay of 0.1 s, number of scans (NS) of 16, and 2048 x 256 complex points for the 1H and 15N dimensions, respectively. Spectra were processed using NMRPipe ^102^ on the NMRbox biomolecular NMR processing platform and analyzed with CcpNmr Analysis 3.2.0^103^ to acquire time-dependent peak volume decay of all the resonances.

### Bio-layer interferometry (BLI) and data analysis

BLI experiments were done on the Pall Fortebio Blitz^TM^ Protein Detection System with Sartorius Octet® Streptavidin (SA) Biosensor (Cat#18-5019) at the Keck Biophysics Facility, a shared resource of the Robert H. Lurie Comprehensive Cancer Center of Northwestern University supported in part by the NCI Cancer Center Support Grant #P30 CA060553. Biotin labeled jR2R3-P301L peptides from GenScript were loaded onto the sensors using 20 μM peptide solution. tRNA samples were prepped in the following step prior to measurement: 100 μM stock solution in ultrapure water thawed from -20C storage, heat shocked at 70C for 3 minutes, reannealing at RT for 10 mins, then serial dilutions were made from 10 μM down to 0.0098 μM. 5 to 7 concentrations were tested for each tRNA sample. The affinity measurement was done with 30 seconds in ultrapure water to measure the baseline, 150 seconds in each tRNA concentration to measure analyte association, and then 120 seconds in ultrapure water to measure analyte dissociation.

For kinetic fitting, BLI association and dissociation traces were fit globally to a 1:1 Langmuir binding model using a custom Python script. For association, the response was modeled as *R*(*t*) = *R_eq_*(1 − *e*^(*k_obs_*\*t)^), where *k_obs_* = *k_on_*[*A*] + *k_off_* and 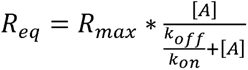. Dissociation was modeled as *R*(*t*) = *R*_0_ * *e*^(− *k_off_*t*)^, where *R_0_* is the response at the end of association. *k_obs_* is the rate representing sensorgram approaching equilibrium during association, *k_on_* is the association rate, *k_off_*, is the dissociation rate, *R_max_* is the capacity, and |*A*| is the analyte concentration. *k_on_* and *k_off_* were treated as global parameters shared across all analyte concentrations, while *R_max_* was fitted locally for each concentration. The equilibrium dissociation constant calculated as 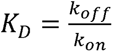 from the kinetic fit is referred to as kinetic *K_D_*, distinct from the value at equilibrium. The uncertainty for kinetic *K_D_* was propagated from the standard errors of *k_on_* and *k_off_* extracted from the covariance matrix of the Jacobian at the solution:

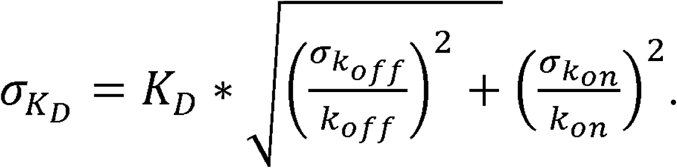

### Flow cytometry analysis

For cell-cycle analysis, cells were collected, washed 3 times with 1X DPBS, fixed, and fixed with 75% ethanol at 20°C overnight. Cells were then further incubated with 5 mM propidium iodide (Sigma) or Vybrant DyeCycle Violet stain (Thermo Fisher Scientific, Cat#V35003) for 30 min at 37°C, followed by the flow cytometry analysis using the BD LSRII analyzer (RRID:SCR_021149). The percentage of sub-G1 was monitored using the FlowJo (RRID:SCR_008520) (v.10.8.1). Annexin V and 7-AAD staining (BD Biosciences, 559763) were performed according to the manufacturer’s instructions. Briefly, the cells were collected by centrifugation, resuspended in 500 mL 1 annexin V binding buffer, incubated with 5 mL annexin V and 5 mL 7-AAD at RT for 15 min in the dark, and followed by the flow cytometry analysis.

### Neurite length and neurite branch point measurement

Live cell imaging was performed using the IncuCyte^TM^ Live-Cell Imaging System (Essen BioScience). Cell confluency, cell body number, neurite length, and branching points were monitored and quantified using the IncuCyte^TM^ software.

### Neuronal viability assays

Rat primary neurons were plated at 40,000 cells/cm^2^ and maintained in 96-well plates for 10 days. Human iNs were plated (15,000 cells/cm^2^) and cultured in 96-well plates for 3 weeks. Following treatments, cell viability was measured using the AlamarBlue HS Cell viability reagent (Life Technologies, Cat#DAL1025) at 1:10 dilution according to the manufacturer’s instructions. After a 4 h incubation at 37 °C, the absorbance of AlamarBlue® was measured at 570 nm, with 600 nm as a reference wavelength using the Promega GloMax 96 Microplate Luminometer (RRID:SCR_018614).

## Data analysis

Data management and calculations were performed using Prism 10 (GraphPad). Comparisons between two groups were performed using the unpaired two-tailed Student’s t-test. For the comparison of more than two groups, one-way analysis of variance (ANOVA), followed by post-hoc test, was performed. For the comparison of more than two independent variables, a two-way ANOVA was performed. P value < 0.05 was considered statistically significant, and the following notations are used in all figures: *P < 0.05, **P < 0.01, ***P < 0.001, and ****P < 0.0001. All error bars shown represent mean +/- standard error of the mean (SEM) unless otherwise stated.

