## Supplementary figures and tables for "tRNA-derived fragments elevated in Alzheimer’s disease promote Tau aggregation"

Ami Kobayashi et al.

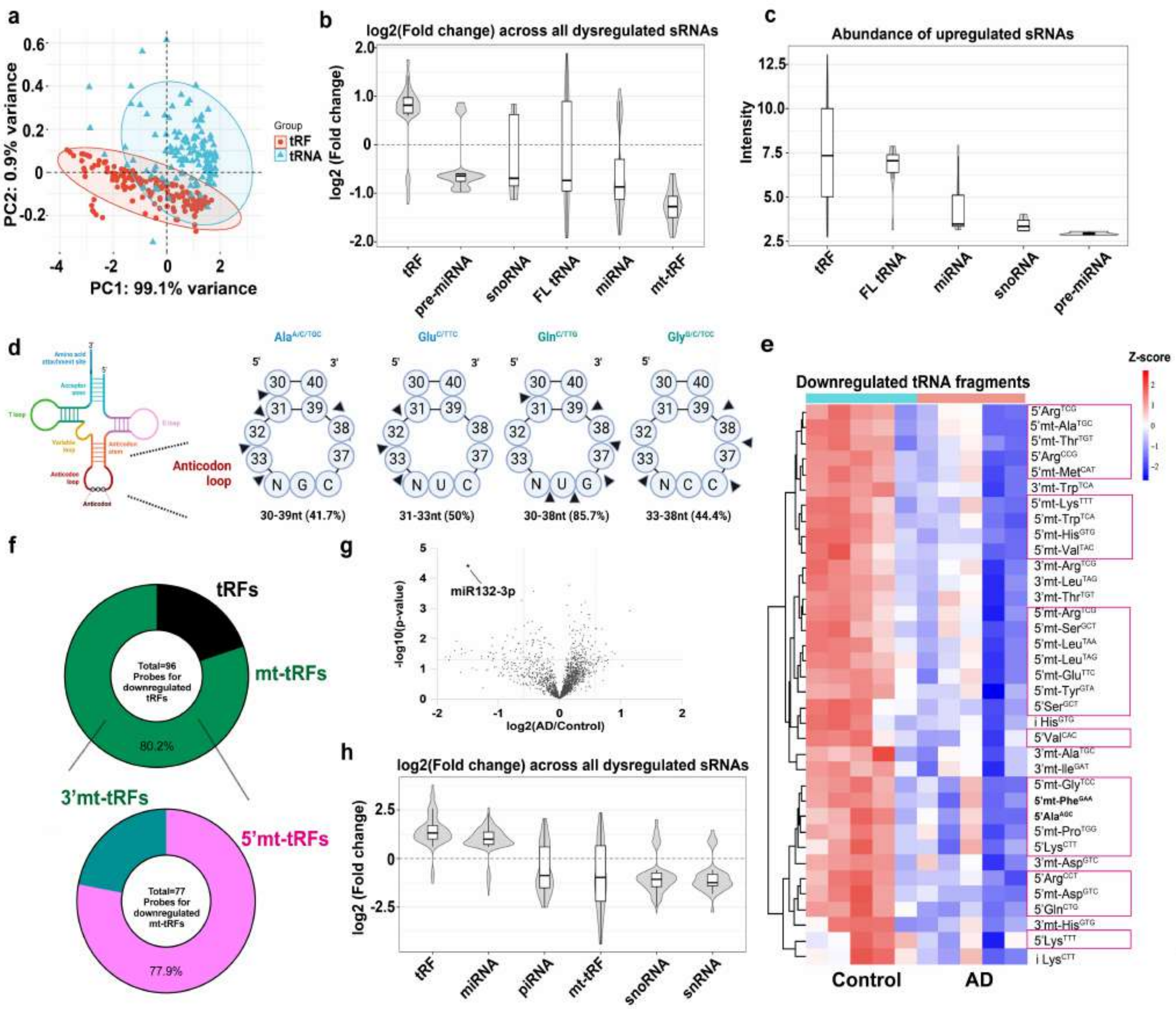


**Supplementary Fig. S1**

**tRFs are dysregulated in human AD patient temporal lobe (related to Figure 1)**

**a.** Principal component analysis (PCA) of the sRNA array data indicates a separation of tRF and fl-tRNA signals. **b**,**c**. Violin plots of log2(Fold Change) across all dysregulated sRNAs (**b**) and abundance of significantly upregulated sRNAs (**c**) from temporal cortex, comparing stage Braak 6 patients (n=5 brain samples per group) to control (Braak 0). **d.** Schematics of tRNA cleavage. Arrowheads indicate cleavage sites, and 5' or 3' end positions of major upregulated tRFs. **e.** A heatmap presenting downregulated tRFs in AD temporal cortex (n=5 per group), with majority corresponding to mt-tRFs. **f**. The proportion of probes detecting mt-tRFs among the downregulated tRFs in AD temporal cortex. The majority of downregulated mt-tRFs are 5'mt-tRFs. **g**. Volcano plot of miRNAs altered in AD temporal cortex, with most downregulated miR-132-3p marked with a black star (n=5 per group). The data are plotted as log2 fold change versus the -log10 of the p-value. Vertical lines: |Fold Change|>1.5, Horizontal lines: p<0.05. **h.** Violin plots of log2(Fold Change) across all dysregulated sRNAs from sRNAseq dataset of prefrontal cortex comparing control (Braak 0 or 1) and AD (stage Braak 5 or 6) (n=6 samples/ group) (GEO accession number GSE48552).

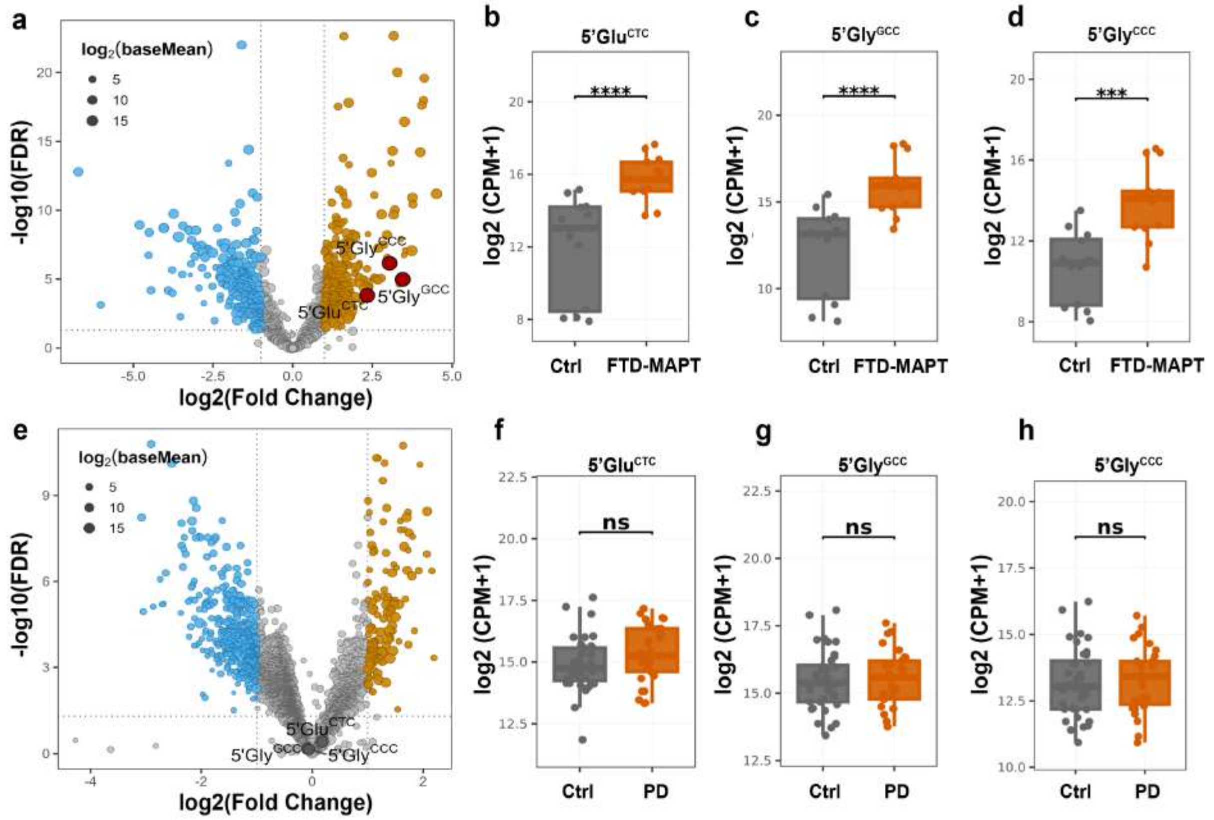


**Supplementary Fig. S2**

**Signature 5′ tRFs are elevated in FTD-MAPT but not in PD**

**a**. Volcano plots of frontal cortex small RNAseq comparing FTD-MAPT (n=13) and control subjects (n=12) with significantly downregulated and upregulated tRFs labeled in blue and brown, respectively. The size of the dots is proportional to the log2 mean expression, and 5'tRFs-5'GluCTC, 5'GlyGCC, and 5'GlyCCC are marked by red circles. Differential expression was evaluated using the DESeq2 Wald test with FDR adjustment. Dataset E-MTAB-12731, batch 1 reanalyzed. **b-d**. Expression of 5'GluCTC, 5'GlyGCC, and5'GlyCCC in FTD-MAPT. Values represent log2(CPM + 1). Boxplots show the median, interquartile range, and individual samples. Group differences were assessed using the Wilcoxon rank-sum test, and significance is indicated above each panel. **e**. Volcano plots of prefrontal cortex small RNAseq comparing PD (n=29) and control subjects (n=36) with significantly downregulated and upregulated tRFs labeled in blue and brown, respectively. The size of the dots is proportional to the log2 mean expression, and 5'tRFs-5'GluCTC, 5'GlyGCC, and 5'GlyCCC are marked by black circles Differential expression was evaluated using the DESeq2 Wald test with FDR adjustment. GEO dataset GSE72962, GSE64977 reanalyzed. **f-h**. Expression of 5'GluCTC, 5'GlyGCC, and5'GlyCCC in PD. Values represent log2(CPM + 1). Boxplots depict median, interquartile range, and individual samples; statistical significance was assessed using the Wilcoxon rank-sum test. Statistically significant differences are indicated as follows: ****p < 0.0001, ***p < 0.001, ns = significant.

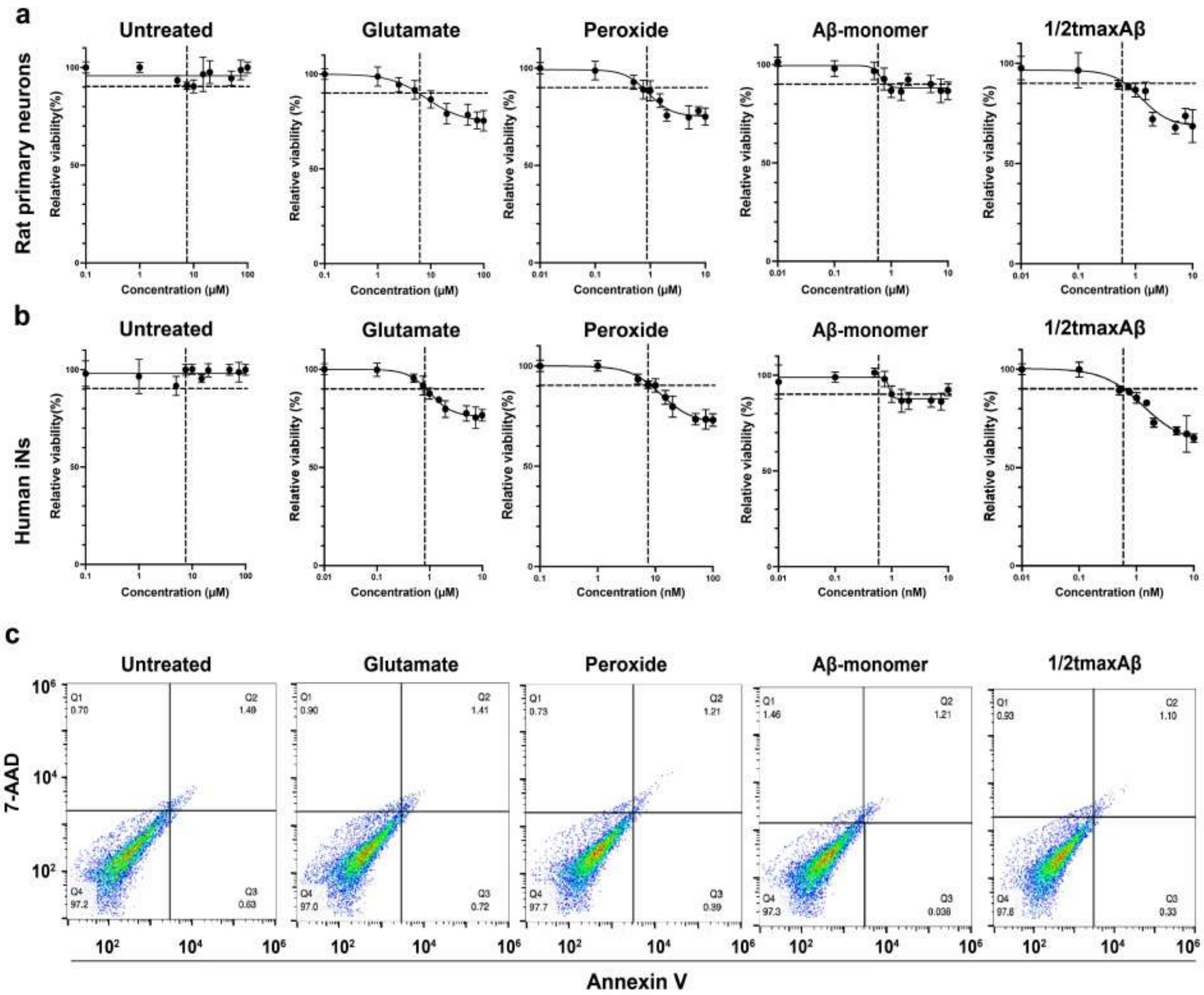


**Supplementary Fig. S3**

**Stress -induced neuronal toxicity is dose-dependent.**

**a**. Dose response experiments with indicated treatments were performed on DIV10 rat primary neurons. Neuronal viability was assessed by AlamarBlue assay 24 hours after treatment (n=3 biological replicates/group). EC90 was calculated using sigmoidal fit. **b**. Dose response experiments with indicated treatments were performed on day 21 human neurons (iNs). Neuronal viability was assessed by AlamarBlue assay 24 hours after treatment (n=3 biological replicates/group). EC90 was calculated using sigmoidal fit. **c.** Rat neurons were treated with 3 µM Glutamate, 0.5 µM Peroxide, 0.25 µM Aβ monomer and 1/2$t_{max}$Aβ for 24 hours, followed by annexin V and 7-AAD staining and flow cytometry analysis (n=3 biological replicates/group).

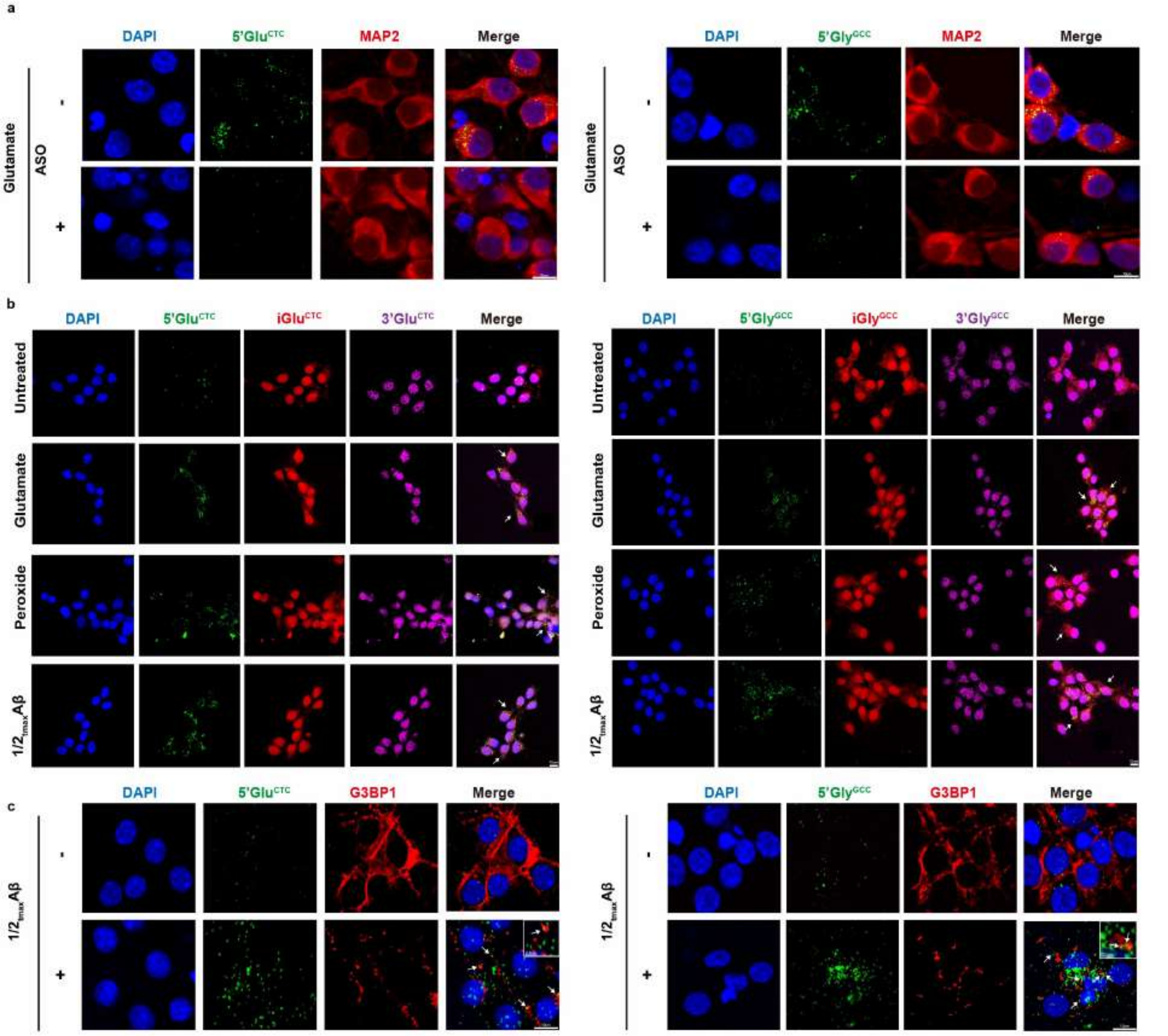


**Supplementary Fig. S4**

**Fluorescent in-situ hybridization (FISH) demonstrates accumulation of 5’Glu$^{CTC}$ and 5’Gly$^{GCC}$.**

**a.** Representative 5’Glu$^{CTC}$ or 5’Gly$^{GCC}$ FISH analyses (green) of glutamate-stressed (3 µM glutamate) rat neurons. Transfection of cognate ASOs 24 hr before cell fixation abolishes the corresponding 5’tRF FISH signals. Scale bar=10µM. **b.** Representative FISH analyses of rat primary neurons, with probes for 5’Glu$^{CTC}$, 5’Gly$^{GCC}$ (green), iGlu$^{CTC}$, iGly$^{GCC}$ (red), and 3’Glu$^{CTC}$, 3’Gly$^{GCC}$ (purple). The cells were treated with 3 µM glutamate, 0.5 µM peroxide, and 0.25 µM 1/2$t_{max}$Aβ for 24 hours (n=3 biological replicates/group). **c.** Representative images of G3BP1 IF (red) with 5’Glu$^{CTC}$ or 5’Gly$^{GCC}$ FISH (green) in rat primary neurons treated with or without 0.25 µM 1/2$t_{max}$Aβ. Arrows indicate colocalization of 5’Glu$^{CTC}$ or 5’Gly$^{GCC}$ with G3BP1. Scale bar=10 µm.

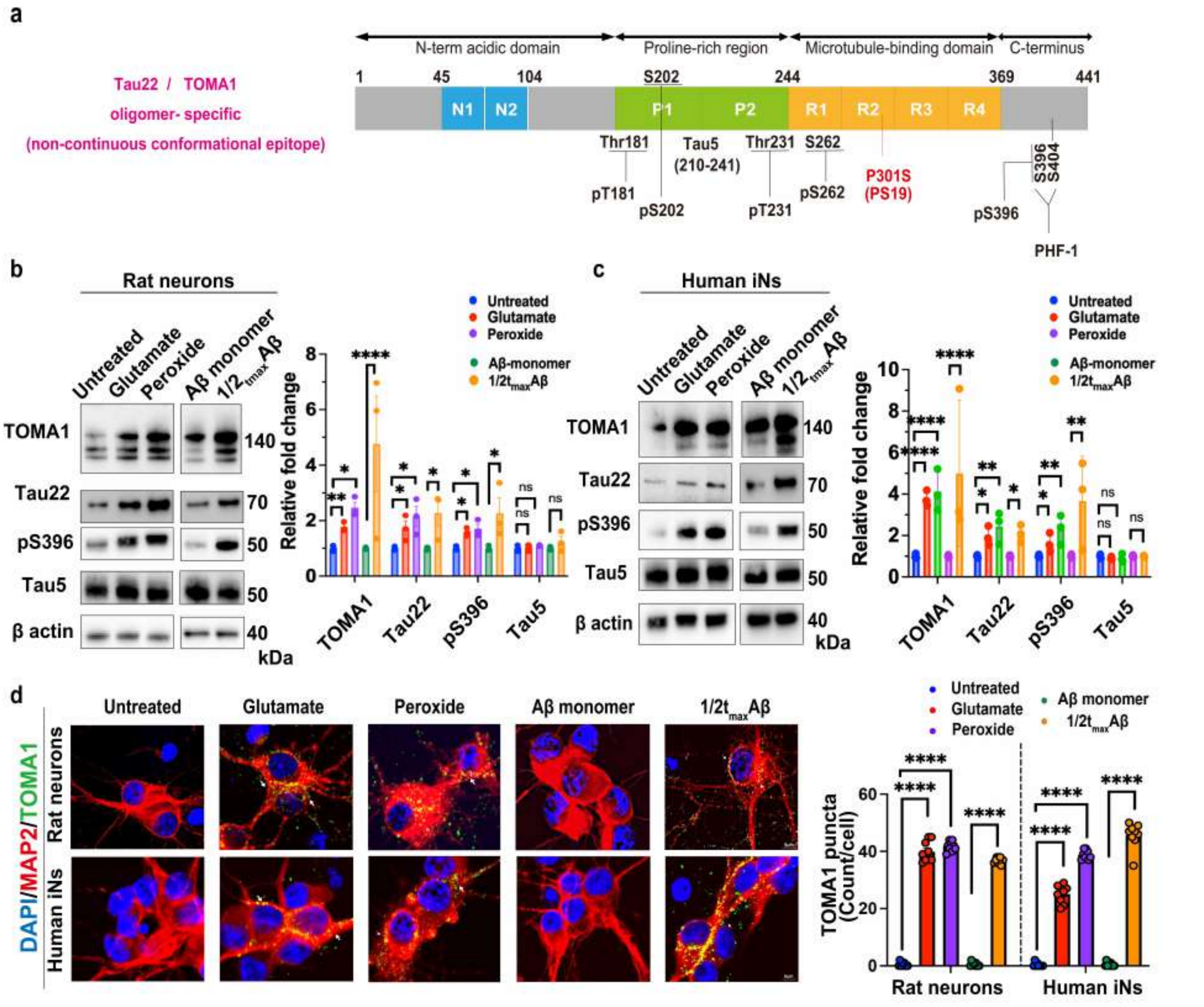


**Supplementary Fig. S5**

**Persistent neuronal stress induces oligomeric and phosphorylated Tau.**

**a.** Schematic presentation of Tau protein domains, with epitopes recognized by the antibodies utilized in this study. **b,c.** Neuronal stresses induce oligomeric Tau (TOMA1, Tau 22) and pTau (Tau pS396) without affecting total Tau (Tau 5) in (**b**) rat primary neurons and (**c**) human iPS-derived neurons (Western blot analysis, one-way ANOVA followed by Dunnett's multiple comparisons test and unpaired two-tailed Student's *t-test* were performed for three or two group comparison, respectively, n=3). **d.** Immunofluorescence for TOMA1 (green) and MAP2 (red) in rat and human neurons, treated as indicated. Rat primary neurons were treated with 3 µM glutamate, 0.5 µM peroxide, and 0.25 µM 1/2$t_{max}$Aβ for 24 hours. Human neurons treated with 0.5 µM glutamate, 5 nM peroxide, and 0.5 nM 1/2$t_{max}$Aβ for 24 hours. Representative images and quantification are shown. At least 200 cells per condition were analyzed across 3 experimental replicates (one-way ANOVA followed by Dunnett's multiple comparisons test and unpaired two-tailed Student's *t-test* were performed for three or two group comparison, respectively, n=9 biological replicates/group). Statistically significant differences are indicated as follows: ****$p < 0.0001$, ***$p < 0.001$, **$p < 0.01$, *$p < 0.05$, ns = not significant.

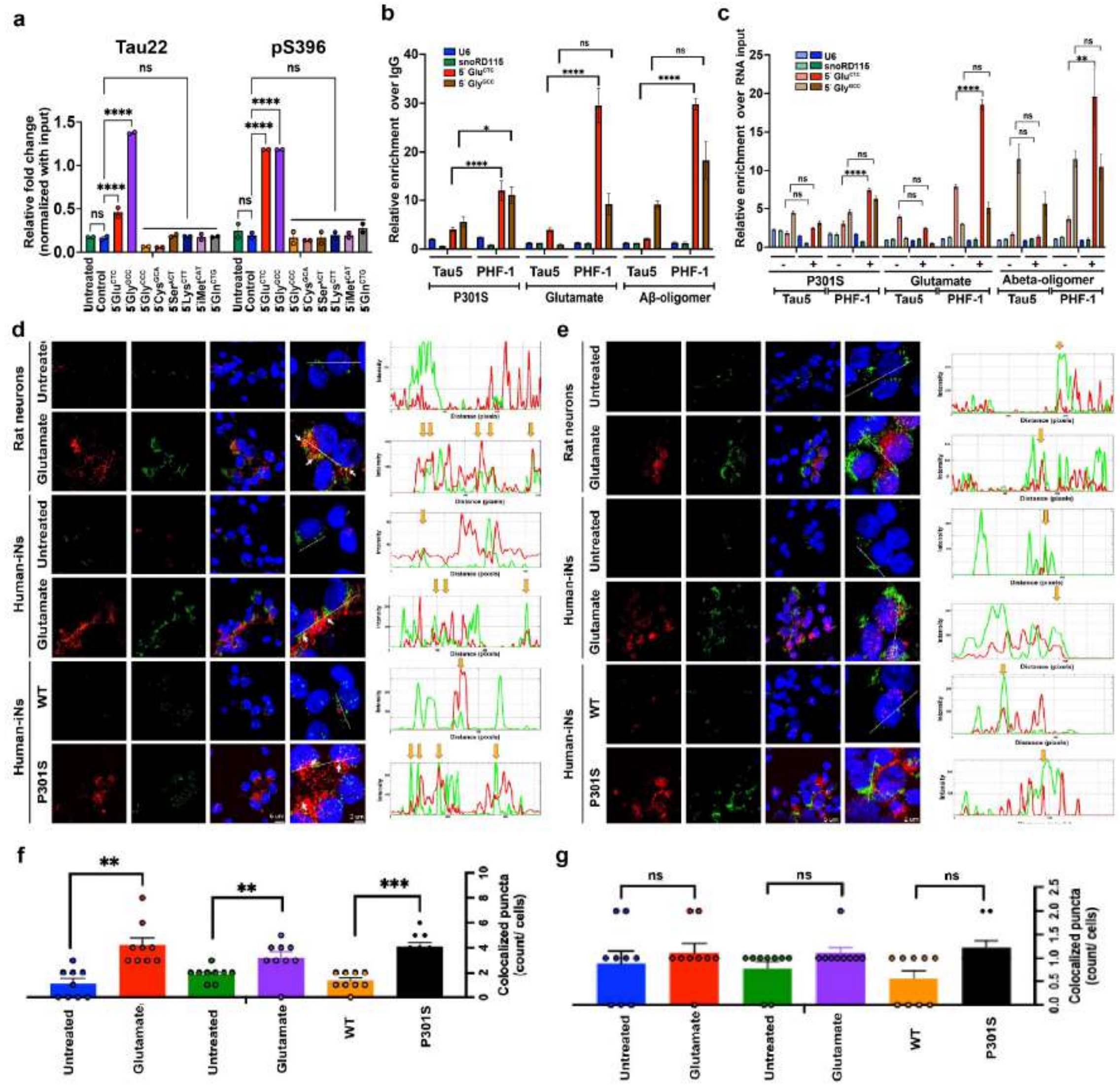

**Supplementary Fig. S6**

**5'Glu$^{CTC}$ is selectively enriched in Tau PHF-1 complexes and colocalizes with oligomeric Tau in neurons**

**a.** Quantification of oligomeric Tau (Tau 22), pTau (Tau pS396) in 5'tRF-bound protein complexes normalized by input demonstrating selective binding of 5'Glu$^{CTC}$ and 5'Gly$^{GCC}$ (n = 2 biological replicates/group). **b,c.** CLIP-qRT-PCR analysis of PS19 (Tau P301S-mut) mouse hippocampi, and rat neurons stressed with either glutamate or 1/2$_{tmax}$Aβ. The bar graph presents the fold enrichment of 5'Glu$^{CTC}$, 5Gly$^{GCC}$, snoRD115, and U6 snRNA in Tau PHF-1 complexes, relative to Tau 5, and normalized to IgG (**b**) and RNA input (**c**) (unpaired two-tailed Student's *t-test*, n = 3 biological replicates). **d,e.** Representative images of TOMA1 IF (red) with (**d**) 5'Glu$^{CTC}$ or (**e**) 5'Gly$^{GCC}$ FISH (green) in rat primary, human WT and Tau P301S mutant iPS-derived neurons. Plot profiles show a spatially resolved graph of fluorescence intensity for each channel. Arrows indicate colocalization of 5'Glu$^{CTC}$ or 5'Gly$^{GCC}$ with TOMA1. **f,g.** Quantification of the colocalized puncta count per cell in **f** (**d**) and **g** (**e**) are shown. At least 150 cells per condition were analyzed across 3 experimental replicates. (unpaired two-tailed Student's t test, n = 9 biological replicates/group).

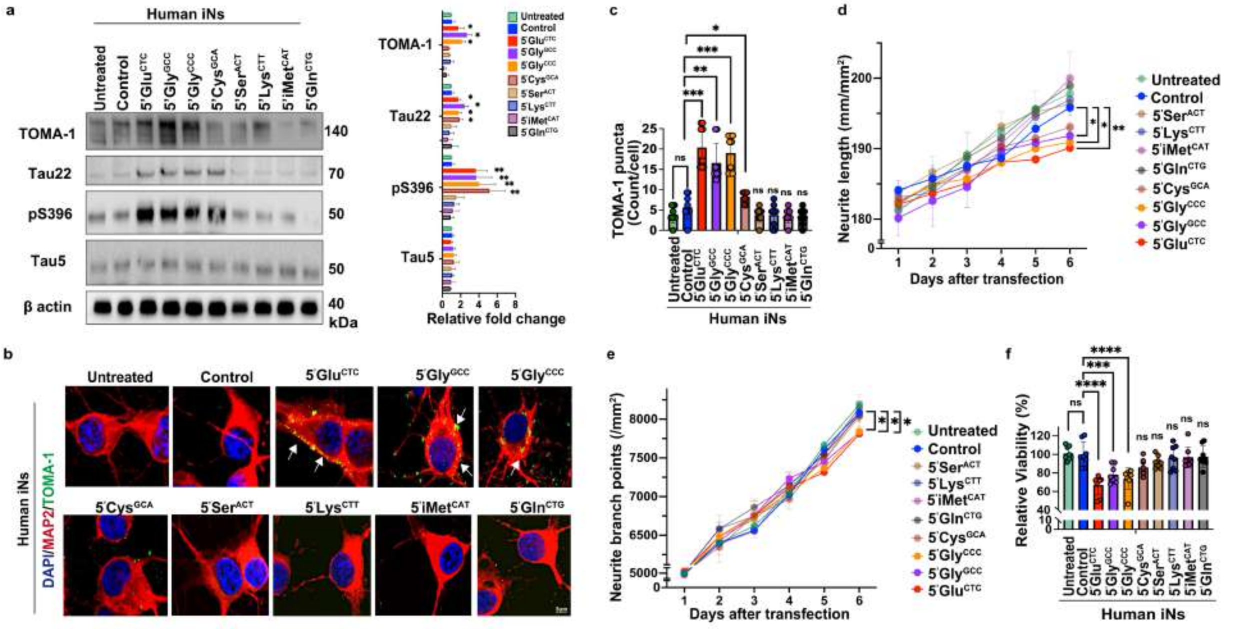


**Supplementary Fig. S7**

**Specific 5'tRFs induce Tau phosphorylation and oligomerization and impair viability of human iPSC-derived neurons.**

**a.** Representative Western blots and quantification of oligomeric Tau (TOMA1, Tau 22), pTau (pS396), and total Tau (Tau 5) in human neurons transfected with indicated 5'tRFs along with control cultures (n = 3 biological replicates/group). **b.** Representative IF images of human neurons transfected with indicated 5'tRFs, showing TOMA1 puncta colocalized with MAP2 (yellow), and nuclei (blue). Arrows mark TOMA1 puncta. **c.** Quantification of TOMA1 puncta in 5'tRF-transfected human neurons. At least 150 cells per condition were analyzed across 3 experimental replicates (one-way ANOVA followed by Dunnett's multiple comparisons test, n = 9 biological replicates/group). **d,e.** Human neurons were transfected with indicated 5'tRFs and monitored for 6 days by live-cell imaging, along with control cultures. Neurite lengths (**d**) and neurite branch points (**e**) were quantified (one-way ANOVA followed by Dunnett's multiple comparisons test, n = 3 biological replicates/group). **f.** Neuronal viability assessed 6 days after 5'tRF-transfection (one-way ANOVA followed by Dunnett's multiple comparisons test, n = 5 biological replicates/group).

All graphs show mean ± S.E.M. ****$p < 0.0001$, ***$p < 0.001$, **$p < 0.01$, *$p < 0.05$, ns = not significant.

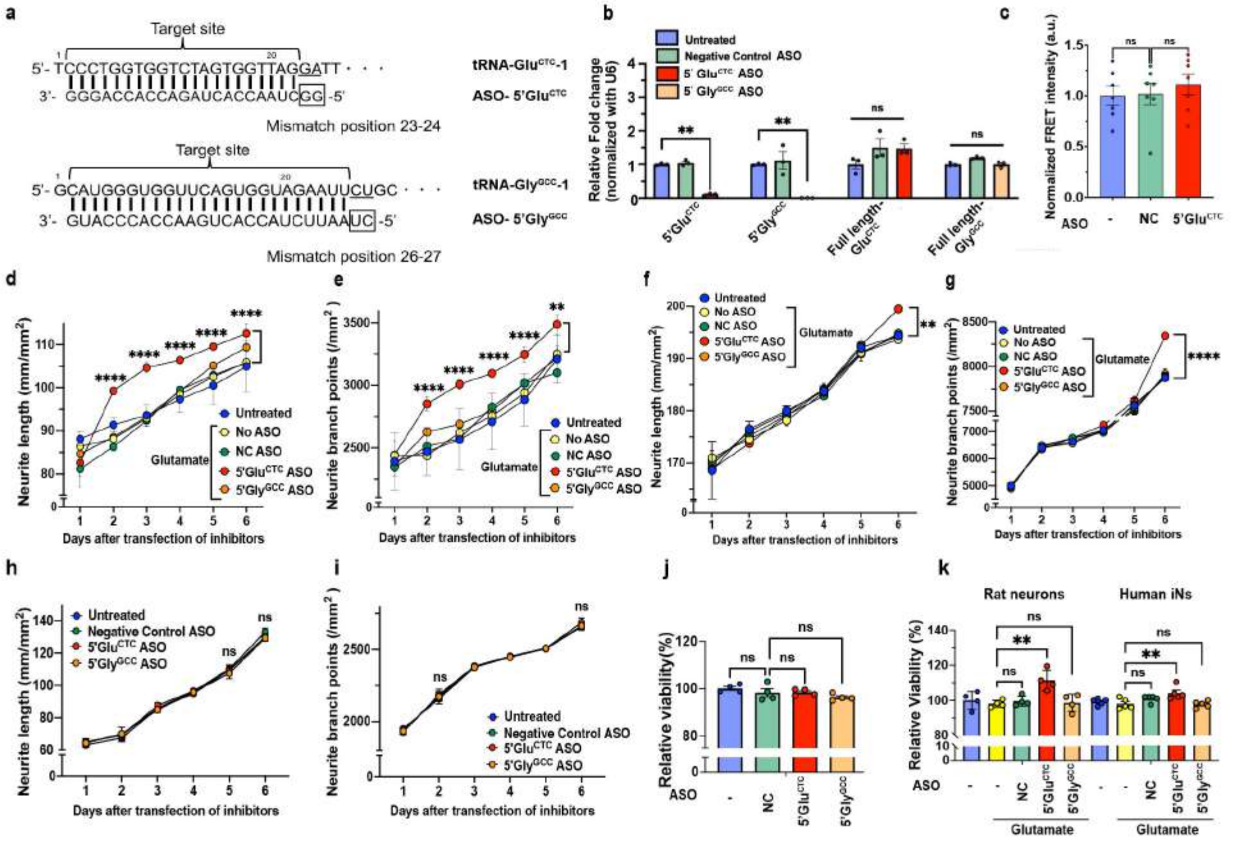


**Supplementary Fig. S8**

**Efficacy, specificity, and effects of 5’Glu$^{CTC}$ and 5’Gly$^{GCC}$ ASOs in rat and human neurons**

**a.** Sequences of 5’Glu$^{CTC}$, 5’Gly$^{GCC}$, and ASO inhibitors. The mismatches within the ASO sequences (depicted by square frames) allow reduced binding to fl-tRNAs and increase the specificity for 5’tRFs. **b.** qRT-PCR analysis of fl-tRNAs and 5’tRFS in neurons transfected with 5’Glu$^{CTC}$ and 5’Gly$^{GCC}$ ASO inhibitors (one-way ANOVA followed by Dunnett’s multiple comparisons test). **c.** Tau seeding assay in HEK-tau-biosensor cells. ASOs for 5’Glu$^{CTC}$ or 5’Gly$^{GCC}$, and HMW tau were co-transfected to the cells, and Tau aggregation quantified by FRET intensity (n = 7 biological replicates/group). **d,e.** Live-cell imaging–based quantification of neurite length (**d**) and neurite branch points (**e**) in glutamate-stressed rat neurons transfected with 5’Glu$^{CTC}$ or 5’Gly$^{GCC}$ ASOs and monitored for 6 days (two-way ANOVA, n = 3 biological replicates/group). **f,g.** Live-cell imaging–based quantification of neurite length (**f**) and neurite branch points (**g**) in glutamate-stressed human iNs transfected with 5’Glu$^{CTC}$ or 5’Gly$^{GCC}$ ASOs and monitored for 6 days (two-way ANOVA, n = 3 biological replicates/group). **h,i.** Live-cell imaging–based quantification of neurite length (**h**) and neurite branch points (**i**) in unstressed rat neurons transfected with 5’Glu$^{CTC}$ or 5’Gly$^{GCC}$ ASOs and monitored for 6 days (two-way ANOVA, n = 3 biological replicates/group). **j.** Neuronal viability assessed 6 days after 5’Glu$^{CTC}$ or 5’Gly$^{GCC}$ ASO transfection in rat neurons (two-way ANOVA, n = 4 biological replicates/group). **k.** Neuronal viability assessed 6 days after 5’Glu$^{CTC}$ or 5’Gly$^{GCC}$ ASO transfection in glutamate-stressed rat neurons and human iNs. (two-way ANOVA compared to untreated controls, n = 3 biological replicates/group).

All graphs show mean ± S.E.M. ****$p < 0.0001$, ***$p < 0.001$, **$p < 0.01$, *$p < 0.05$, ns = not significant.

**Supplementary Table S1**

**a.**

| Cohort | Age | Gender | PMI (hrs) | Brain Region | Diagnosis |
|---|---|---|---|---|---|
| New York Brain Bank at Columbia University | 77 ± 3.7 | 3M/3F | 9.9 ± 6.6 | Temporal cortex | NDAR/Control |
| | 85.3 ± 6.4 | 2M/1F | 20.9 ± 1.4 | Temporal cortex | Braak III |
| | 82.8 ± 5.3 | 1M/5F | 14.2 ± 7.9 | Temporal cortex | Braak VI |
| Mount Sinai Brain Bank (MSBB) | 84.6 ± 2.1 | 5M/5F | 5.4 ± 1.5 | Temporal cortex | NDAR/Control |
| | 82.5 ± 1.1 | 6M/6F | 3.7 ± 1.4 | Temporal cortex | Braak II-VI |
| Masachusetts Alzheimer's Disease Research Center (ADRC) | 84 | M | 7 | Frontal cortex | Braak VI |

Data presented as mean with SEM.
M: males, F: females, PMI: Post Mortem Interval, NDAR: No Diagnostic Abnormality Recognized
All data are mean ± standard deviation

**b.**

| Cohort | Age | Gender | PMI (hrs) | Brain Region | Diagnosis |
|---|---|---|---|---|---|
| New York Brain Bank at Columbia University | 72 | M | 5.7 | Temporal cortex | NDAR/Control |
| | 82 | F | 22.8 | Temporal cortex | NDAR/Control |
| | 78 | M | 8 | Temporal cortex | NDAR/Control |
| | 76 | F | 5.4 | Temporal cortex | NDAR/Control |
| | 80 | F | 6.8 | Temporal cortex | NDAR/Control |
| | 74 | M | 10.9 | Temporal cortex | NDAR/Control |
| | 89 | M | 20.3 | Temporal cortex | Braak III |
| | 89 | F | 20.1 | Temporal cortex | Braak III |
| | 78 | M | 22.6 | Temporal cortex | Braak III |
| | 89 | F | 15.2 | Temporal cortex | Braak VI |
| | 78 | F | 27.6 | Temporal cortex | Braak VI |
| | 75 | M | 6.2 | Temporal cortex | Braak VI |
| | 85 | F | 17.1 | Temporal cortex | Braak VI |
| | 86 | F | 6.5 | Temporal cortex | Braak VI |
| | 84 | F | 12.6 | Temporal cortex | Braak VI |
| Mount Sinai Brain Bank (MSBB) | 85 | F | 5.3 | Temporal cortex | NDAR/Control |
| | 86 | F | 4.4 | Temporal cortex | NDAR/Control |
| | 88 | M | 8 | Temporal cortex | NDAR/Control |
| | 85 | F | 5 | Temporal cortex | NDAR/Control |
| | 85 | F | 4.3 | Temporal cortex | NDAR/Control |
| | 80 | F | 4.8 | Temporal cortex | NDAR/Control |
| | 86 | M | 7.9 | Temporal cortex | NDAR/Control |
| | 84 | M | 2.9 | Temporal cortex | NDAR/Control |
| | 85 | M | 5.3 | Temporal cortex | NDAR/Control |
| | 82 | M | 6.3 | Temporal cortex | NDAR/Control |
| | 84 | F | 2 | Temporal cortex | Braak VI |
| | 84 | F | 2.8 | Temporal cortex | Braak VI |
| | 84 | F | 2.6 | Temporal cortex | Braak V |
| | 81 | M | 4.5 | Temporal cortex | Braak VI |
| | 81 | F | 2.4 | Temporal cortex | Braak III |
| | 83 | F | 2.4 | Temporal cortex | Braak VI |
| | 83 | F | 2.1 | Temporal cortex | Braak VI |
| | 81 | M | 5.6 | Temporal cortex | Braak II |
| | 83 | M | 4.8 | Temporal cortex | Braak II |
| | 82 | M | 4.3 | Temporal cortex | Braak IV |
| | 82 | M | 5.8 | Temporal cortex | Braak VI |
| | 82 | M | 5.5 | Temporal cortex | Braak VI |

M: males, F: females, PMI: Post Mortem Interval, NDAR: No Diagnostic Abnormality Recognized

**Supplementary Table S2**

| Oligos | Sequences |
|---|---|
| 5’Glu$^{CTC}$ | UCC CUG GUG GUC UAG UGG UUA GGA UUC GGC G |
| 5’Gly$^{GCC}$ | GCA UUG GUG GUU CAG UGG UAG AAU UCU CGC C |
| 5’Gly$^{CCC}$ | GCG CCG CUG GUG UAG UGG UAU CAU GCA AGA U |
| 5’Cys$^{GCA}$ | GGG GGU AUA GCU CAU UGG UAG AGC AUU UGA |
| 5’Ser$^{GCT}$ | GUA GUC GUG GCC GAG UGG UUA AGG CGA UGG |
| 5'Lys$^{CTT}$ | GCC CGG CUA GCU CAG UCG GUA GAG CAU GGG ACU |
| 5'iMet$^{CAT}$ | AGC AGA GUG GCG CAG CGG AAG CGU GCU GG |
| 5'Gln$^{CTG}$ | GGU UCC AUG GUG UAA UGG UUA GCA CUC UG |
| Control | UGU GAG UCA CGU GAG GGC AGA AUC UGC UC |

Sequences of synthetic 5’tRFs used in this study.

**Supplementary Table S3**

| Oligo | Sequences |
|---|---|
| 5'Glu$^{CTC}$ inhibitor | mC/ZEN/mCmUmAmAmCmCmAmCmUmAmGmAmCmCmA<br>mCmCmAmGmGmG/3ZEN/ |
| 5'Gly$^{GCC}$ inhibitor | mG/ZEN/mAmAmUmUmCmUmAmCmCmAmCmUmGmAmA<br>mCmCmAmCmCmCmAmUmG/3ZEN/ |
| Negative Control inhibitor | mG/ZEN/mCmGmAmCmUmAmUmAmCmGmCmGmCmAmA<br>mUmAmUmGmG/3ZEN/ |

The sequence of synthetic 5’tRF-inhibitors used in this study.
mA/mU/mC/mG=2’O-methyl (2’OMe) RNA base; /ZEN/ or /3ZEN/=internal or 3’ZEN modification.
All oligonucleotides used in this study were synthesized and purifies by Integrated DNA Technology.
All oligonucleotides are at least 95% homogenous.

**Supplementary Table S4**

| qRT-PCR primers | Sequences (5' to 3') |
|---|---|
| 5'Glu$^{CTC}$ | F: TCCCTGGTGGTCTAGTG |
| | R: CTGCGATGAGTGGCAGGC |
| 5'Glu$^{TTC}$ | F: TCCCATATGGTCTAGCGG |
| | R: CTGCGATGAGTGGCAGGC |
| 5'Gly$^{GCC}$ | F: GCATTGGTGGTTCAGTG |
| | R: CTGCGATGAGTGGCAGGC |
| 5'Gly$^{CCC}$ | F: GCGCCGCTGGTGTAGTGG |
| | R: CTGCGATGAGTGGCAGGC |
| 5'Cys$^{GCA}$ | F: AGTGGTAGAGCATTTGACTGC |
| | R: CTGCGATGAGTGGCAGGC |
| 5'Ala$^{TGC}$ | F: GGGGATGTAGCTCAGTGG |
| | R: CTGCGATGAGTGGCAGGC |
| Full-length-Glu$^{CTC}$ | F: CCTGGTGGTCTAGTGGTTAGG |
| | R: TCCCTGACCGGGAATCGAA |
| Full-length-Gly$^{GCC}$ | F: GCATTGGTGGTTCAGTGGTAGA |
| | R: GCATTGGCCGGGAATCGAA |

The primer sequences of qRT-PCR for 5'tRFs and full-length tRNAs used in this study. F, Forward; R, Reverse.

**Supplementary Table S5**

| **Northern blot Probes** | **Sequences** |
|---|---|
| tRNA 5'Glu$^{CTC}$ (position 1-27) | 5'-GAATCCTAACCACTAGACCACCAGGGA-3' |
| tRNA 5'Gly$^{GCC}$ (position 1-21) | 5'-CTACCACTGAACCACCCATGC-3' |

The sequence of DNA oligo probes for Northern blotting used in this study.

| **FISH Probes** | **Sequences** |
|---|---|
| tRNA 5'Glu$^{CTC}$ (position 2-31) | 5’-CGCCGAATCCTAACCACTAGACCACCAGGG-3’ |
| tRNA i Glu$^{CTC}$ (position 31-52) | 5’-CCCGGGCCGCGGCGGTGAGAGC-3’ |
| tRNA 3'Glu$^{CTC}$ (position 48-70) | 5’-CCCTGACCGGGAATCGAACCCGG-3' |
| tRNA 5'Gly$^{GCC}$ (position 1-31) | 5’-GGCGAGAATTCTACCACTGAACCACCCATGC-3’ |
| tRNA i Gly$^{GCC}$ (position 34-55) | 5’-CGAACCCGGGCCTCCCGCGTGG-3’ |
| tRNA 3'Gly$^{GCC}$ (position 50-69) | 5’-GCATTGGCCGGGAATCGAACC-3' |

The sequence of probes for FISH used in this study.
